# Calbindin-containing CA1 pyramidal cells support cognitive flexibility in spatial task in mice

**DOI:** 10.64898/2025.11.30.691413

**Authors:** Anne Voigt, Verjinia D. Metodieva, Denis Alevi, John J. Tukker, Alexander Stumpf, Daniel Parthier, Dietmar Schmitz

## Abstract

The hippocampus, and particularly the dorsal CA1, is essential for spatial memory consolidation through sharp-wave ripple events (SWRs) that enable the transfer of information to cortical areas. Within the dorsal CA1, distinct pyramidal cell sub-populations of the deep and superficial layer may play distinct roles in the processing and updating of spatial information. Using a Cre-dependent mouseline, we were able to precisely target the superficial, Calbindin (CALB1+) PCs of the CA1. This allowed us to gain a deeper understanding of the connectivity of CALB1+ CA1 PCs to their SWR-propagating excitatory output partners within the Subiculum (SUB), the burst-firing SUB PCs and their functional relevance in spatial memory consolidation and recall. Retrograde rabies tracing revealed heterogeneous innervation of the VGlut2+ bursting PCs by both CA1 PC sub-layers, and showed that the majority of presynaptic inputs are located in the superficial CA1 PC layer. We were able to observe that this anatomically confirmed connection between CALB1+ CA1 PCs and both SUB PC subtypes is able to induce spiking more reliably in burst- than regular-firing SUB PCs. CNO-induced inhibition in a Barnes Maze task revealed that the experimental group showed reduced cognitive flexibility and were slower to adapt to re-location of the goal when CALB1+ CA1 PCs were inhibited during the recall (test), while both groups behaved similarly when consolidation was manipulated (training). Inhibition did not impact overall learning, strategy development or locomotor control. This suggests that CALB1+ CA1 PCs preferentially connect to bursting SUB PCs, and support cognitive flexibility needed to adapt to a changing environment, adding further proof to the functional relevance of laminar segregation of the CA1 and the hippocampus in spatial memory processes.

## Introduction

We have grown tired of the old system – and set out to rearrange our office one day. Pens go where the books once went, and the following days, we keep reaching for the wrong drawer: a reflection of the stability of the spatial map of our surroundings, and a testament to the effort required for new information to out compete older, integrated and consolidated memories. The encoding of memories relies on the repeated activation of neuronal ensembles that previously fired together (Hebb, 1949; Brown, 2020; Cutsuridis et al., 2010; Nadel and Maurer, 2020), a process achieved physiologically through sharp-wave ripple events (SWRs), network oscillation events crucially involved in the consolidation of memories and the transfer of activity from hippocampal to cortical areas (Böhm et al., 2015). As was shown by O’Keefe in 1971, the hippocampus is highly involved in spatial learning and memory functions (O’Keefe and Dostrovsky, 1971). By recording and observing changes in cellular action potentials, they were able to show that the hippocampus is deeply involved in the creation and storage of a “cognitive (spatial) map”, as cells fired at specific locations as the animals moved through space (O’Keefe, 1976; Tolman, 1948). This work cemented the role of the hippocampus in encoding both spatial and non-spatial forms of declarative memory (Basu and Siegelbaum, 2015; Colgin, 2020; Lisman et al., 2017; O’Keefe and Nadel, 1978).

Since then, many studies have shown the functional segregation of the hippocampus along three orthogonal axes, as well as its intrinsic organization (transverse, radial, longitudinal: dorso-ventral axis), indicating a need to understand the hippocampus and its subareas not as a functional monolith, but as a heterogeneous sum of distinct, functionally specialized parts (DG, CA3, CA1, SUB). Without an acknowledgment of this heterogeneity, we risk misinterpretations and inconsistencies in our understanding of the hippocampus (Henriksen et al., 2010). This was first clearly observed after lesions of the dorsal hippocampus gradually blocked learning in a spatial navigation task, whereas lesions of similar and larger scale of the ventral hippocampus did not (Moser et al., 1993, 1995), and has been supported since by studies diving deeper into each subarea, taking cellular composition, orientation and connectivity into account when describing a subarea’s functional relevance in spatial learning. These detailed studies have allowed us to understand the dorsal CA1 as one of the key areas required for spatial information processing and encoding (Hunsaker et al., 2008; Kesner et al., 2010; Lee et al., 2005; Park et al., 2021). Much like the hippocampus itself, function and cellular composition of the CA1 is also heterogeneous. Along the dorsal-ventral axis, robust gene markers and physiological differences separate dorsal CA1 from ventral CA1, with the dorsal region supporting spatial processing and encoding more reliably than the ventral (Cembrowski et al., 2016; Dong et al., 2009; Fanselow and Dong, 2010; Jimenez et al., 2018; Thompson et al., 2008). Electrophysiological recordings revealed pyramidal cells (PCs) in the dorsal CA1 to exhibit more place cells overall, firing at a higher spatial selectivity, resulting in a higher resolution of space per cell (Jung et al., 1994; Jimenez et al., 2018). Dorsal CA1 PCs can also be modulated by environmental context (Harland et al., 2021) and encode the spatial location of rewards via increased activity (Jarzebowski et al., 2022). Temporal deactivation of these cells using CNO in a dynamic foraging task reduced the learning rate of mice in reward-based tasks (Jeong et al., 2018), as the animals were slower to adapt to new reward situations (Tran et al., 2008; Dillon et al., 2008).

### Calbindin marks functionally distinct PC subpopulation on the radial axis of the CA1

Within the circuitry of the hippocampal formation, the dorsal CA1 has also been implicated in maintaining overall sparse neural activation as a key regulator of hippocampal oscillations. Relevant for exploration and spatial learning (Buzsáki, 2002; Winson, 1978), theta oscillations (4–10 Hz in rodents) are markers of organized neural firing (Jacobs, 2014) and prominent in the CA1. Any changes in the alignment of spikes with the troughs of theta waves upon silencing cells in the dorsal CA1 would indicate a loss in temporal control and specificity of hippocampal oscillations, necessary for memory consolidation and the relay of information from the hippocampus to its downstream cortical targets (Adaikkan et al., 2024). Not all cells found in the dorsal CA1 contribute to its broad functional relevance equally: even on a cellular level, we can again observe functional segregation, based on distinct electrophysiological, genetic, morphological, developmental and molecular properties (Soltesz and Losonczy, 2018). Historically, pyramidal cells along the radial axis of the CA1 were segregated by their soma position (deep layer towards the stratum oriens, superficial layer closer to stratum radiatum) (Lorente De Nó, 1934). Using soma location to understand the functional relevance of the pyramidal cells along the deep and superficial layers of the CA1 has revealed that though all cells may be excitatory pyramidal cells, they, too, are not a homogeneous monolith: superficial and deep PCs each display distinct firing properties and fire at different points of theta oscillations, indicating network-state dependent functional segregation (Mizuseki et al., 2011; Danielson et al., 2016). PCs of the superficial layers display greater stability of place maps and manifolds (Esparza et al., 2025; Valero et al., 2015; Sharif et al., 2021), properties which might be related to a distinct molecular marker of this layer: CALB1, a calcium-binding protein, is expressed primarily in sPCs (Baimbridge et al., 1991) and is only one of many markers defining the pyramidal cells of the superficial layer. Calcium signals in neurons are, from a biophysical perspective, short lived (Faas et al., 2011), and cells expressing CALB1 exhibit a biophysical feature needed to precisely contain and constrain local changes in calcium concentration, affecting synaptic integration, plasticity thresholds and spike timing of those cells (Eggermann et al., 2011; Kim et al., 2021; Müller et al., 2005). This is reflected in observations of distinct modulations of CALB1+ sPCs by ripples (Mizuseki et al., 2011; Gu et al., 2023).

### The importance of subtype-specific microcircuits in information processing and relay: Subicular bursting PCs as gateway to the cortex

The functional segregation of the hippocampus and its subfields may not only be due to the components and properties they exhibit. Orientation and location of cell populations is also critically defining for input and output connectivity, and their presence in specific microcircuits. Much like we have just seen for the CA1, the Subiculum (SUB), the output hub of the hippocampus, also contains two principal cell populations, which can be distinguished by their intrinsic firing properties, location along the transverse axis of the hippocampus and genetic marker: burst- and regular-firing pyramidal cells (Wozny et al., 2008; Böhm et al., 2015, 2018; Cembrowski and Menon, 2018). These populations also differ in their output projections, which may support distinct functional roles in memory consolidation and recall. Bursting SUB PCs, which express VG-lut2 (Wozny et al., 2018), have already been shown to play a distinct role in memory consolidation, potentially through sharp-wave ripple (SWR) propagation. Nitzan et al. (2020) demonstrated that SWRs propagate from bursting SUB cells to retrosplenial cortex, carrying hippocampal information to the granular retrosplenial cortex (gRSC), a part of a cortical network (Nitzan et al., 2020). Cembrowski et al. (2018) showed that optogenetic inhibition of the distal SUB, which predominantly contains bursting SUB PCs, was able to impair spatial learning in a Morris water maze, while inhibition of the proximal SUB had no effect (Cembrowski et al., 2018). Together, these findings suggest that bursting SUB PCs may serve to support spatial learning and memory Ding et al. (2022).

Given the importance of cellular subpopulations, their distinct genetic markers and unique distribution along different axes, we must ask: does the laminar organization of CA1 map onto its direct hippocampal output, the SUB and its respective cellular subpopulations? It has already been shown that superficial CALB1+ CA1 PCs are able to maintain stable representations of spatial landmarks that persist when environmental cues change (Danielson et al., 2016; Mizuseki et al., 2011; Geiller et al., 2017; Sharif et al., 2021; Esparza et al., 2025). Does this segregation of the CA1 extend to preferential connectivity with specific SUB populations, and does it give these connections functional relevance? Answering these questions is essential to understand how hippocampal microcircuits, and their role in the computation and translation of information into the cortical areas, support memory consolidation and recall.

## Materials and Methods

All experiments were conducted in accordance with European guidelines and with permission from local regulatory authorities (Berlin Landesamt für Gesundheit und Soziales, permits: G0092/15, G0030/20, G0228/20, T0100/03, G0102/18) and conformed to all relevant regulatory standards.

### Animal Husbandry

Animals used were naive CALB1-Cre, CALB1-Cre × Ai9, VGlut2-Cre (JAX: 016963) and wildtype animals (C57BL/6). The mice were group housed within the in-house animal facility, and kept on a 12-h light/12-h dark cycle at a constant temperature (23-24 °C) and relative humidity (40-50%). Food and water were provided ad libitum.

### Stereotactic injections and viral tools

The mice were anesthetized using a isofluorane (2% vol/vol in oxygen). Before any further surgical procedures, they were given the analgesics carprofen and subcutaneous lidocaine at the incision site. Following this, the mice were positioned in a stereotaxic frame and secured with ear bars. The animals were kept warm with a heating pad placed underneath the stereotactic frame and their body temperature kept at 37°C and their eyes moisturized with lotion during the entire procedure. The skull was levelled and the injection target was located using a micromanipulator (Neurostar, Tübingen, Germany). Virus were injected using a beveled NanoFil syringe (World Precision Instruments, Sarasota County, FL, US). Coordinates used for craniotomies and injections differed depending on the target region. SUB: −3.0 AP; ±1.6 ML; −1.6 DV (Nitzan et al., 2020), used for rabies tracing. CA1: −2.2 AP; ±1.8 ML; −1.5 DV used for injection of optogenetic tools and Barnes Maze experiments. Animals were given at least 2 weeks to recover before the start of the Barnes Maze task. For rabies tracing, animals were kept in their home cages for exactly 3 weeks before the second (rabies) virus was applied, and then terminated after exactly 10 days. The reliability of injections was tested using Cre-dependent GFP-expression in the soma of cells (AAV-DIO-GFP, BA002h, CCO Viral Core Facility Berlin (VCF) https://vcf.charite.de/en/catalog) in CALB1-Cre mice and incubated for 2 weeks. Slice electrophysiology experiments were conducted after at least 3 weeks of virus incubation time. Injections were conducted as described under stereotactic injections. Viral injections for electrophysiological experiments included a Cre-dependent Channelrhodopsin (ChR2) GFP-expressing virus (pAAV-EF1a-Switch-mRubyNLS-hChR2(H134R)-EYFP-NO-WPRE, #362b, VCF). Electrophysiological control experiments used a general ChR2-expressing virus (pAAV-hSyn-hChR2(H134R)-eYFP-WPRE, BA006, VCF) in CALB1-Cre mice. Retrograde rabies tracing used a two-virus system in VGlut2-Cre mice: a Cre-dependent AAV starter virus at 200nl, 50nl/min as starter virus (AAV1-Syn-Flex-nGToG-WPRE3, BA96d, VCF) expressing TVA and RG in glutamatergic cells, and a glycoprotein-deleted rabies virus CVS-N2C virus (RABV CVS-N2c(deltaG)-DsRed), BRVenvA-24, VCF) for monosynaptic retrograde labeling. Mice were kept in home cages for 3 weeks after starter virus injection before rabies virus application, followed by a 10-day incubation. Behavioral experiments used 500nl, at 40nl/min DREADDs (pAAV-hSyn-DIO-hM4D(Gi)-mCherry, #333b, VCF) incubation time for 3 weeks.

### Brain Sectioning using BrainSaw

Injections followed description of stereotactic injection. For decapitation, animals were anesthetized with 4-5% isoflurane in O_2_; brains were removed and placed in 4°C cold 4% PFA. For perfusions, deep anesthesia was induced using urethane (2.5 g/kg BW), followed by transcardial perfusion with PBS and 4% PFA. All brains were post-fixed overnight at 4°C in PFA and transferred to PBS. Brains were embedded in 4% agarose and sliced using a custom-built two-photon serial sectioning microscope. Imaging was performed using ScanImage (Vidrio Technologies, USA) and BakingTray (https://bakingtray.mouse.vision), both MATLAB-based. The setup consisted of a 2P microscope coupled to a vibratome (VT1000S, LEICA) and a high-precision motorized stage (X/Y: V-580; Z: L-310, Physik Instrumente). Imaging was done with a 20× water-immersion objective at 0.8 *µ*m XY resolution and 5 *µ*m axial PSF. Fluorescence was collected in the green channel (500-550 nm, ET525/50). Each brain was imaged in 1,025 × 1,025 *µ*m tiles with 7% overlap. Final voxel size: 0.8 × 0.8 × 5 *µ*m (XYZ). Physical slicing thickness was set at 50 *µ*m, with optical sectioning at 5 *µ*m using a piezo objective scanner (PIFOC P-726, Physik Instrumente). Dual-channel acquisition included the red channel (580-630 nm, ET605/70, Chroma). Tile stitching was performed using StitchIt (https://github.com/SWC-Advanced-Microscopy/StitchIt), which corrected illumination and combined tiles. If multiple brains were acquired in one run, StitchIt.SampleSplitter was used to separate and label datasets. Data analysis was performed using napari, BrainReg, and custom Python scripts.

### Two-Photon serial tomography

After PBS-wash, the brains were embedded in 4% agarose, then sliced and imaged using a custom-made two-photon serial sectioning microscope. The procedure was run using ScanImage (Vidrio Technologies, USA) and BakingTray (https://bakingtray.mouse.vision), both MATLAB-based software tools. ScanImage is used to perform imagine, and BakingTray an extension to slice a sample after each sectioning. The entire setup consists of a two-photon microscope paired with a vibratome (VT1000S, LEICA, Germany) and a X/Y/Z high precision stage (X/Y: V-580; Z: L-310, Physik Instrumente, Germany). A 20× water immersion objective was used with a resolution of 0.8 *µ*m in X and Y and measured axial point spread function (PSF) at ∼5 *µ*m full width at half maximum. We collected fluorescence in the green channel (500-550 nm, ET525/50) and each section consisted of 1,025 × 1,025 *µ*m tiles overlapping at 7 percent. The final voxel size was 0.8 × 0.8 × 5 *µ*m (X, Y, Z). The brains were sliced at 40*µ*m thickness. The thickness of physical slices was set to be 50 *µ*m for the entire brain and we acquired optical sections at 5 *µ*m using a highly dynamic piezo objective scanner (PIFOC P-726, Physik Instrumente, Germany) in two channels (green channel: 500-550 nm, ET525/50, Chroma, USA; red channel: 580-630 nm, ET605/70, Chroma, USA). Each brain section was imaged with 7% overlapping 1025 × 1025 *µ*m tiles. The raw tiles were stitched using another MATLAB-based software and companion package to BakingTray, StitchIt (https://github.com/SWC-Advanced-Microscopy/StitchIt). StitchIt combines the tiles based on the position of the tile position and corrects the illumination based on the average tile of each optical plane. When two brains were sliced and imaged at the same time, this step was followed by using StitchIt. Sample-Splitter, an extension of StitchIt, which allows for the user to crop, label and split up an acquisition of multiple samples, and separate them into distinct repositories.

### Cell Detection and Processing

Quantification of cells was achieved by registering the sliced and imaged brain sections (whole brain) to the Allen Mouse Brain Atlas using Brain-Reg, followed by manual and automated cell detection using CellFinder of pre-synaptic cells within the CA1. Data sets were excluded when injection coordinates and viral spread were off-target. Following cell detection, spatial distribution of labeled PCs in the CA1 were analyzed to determine their positions relative to the SP/SR border (strpyr line). This line was drawn manually every 5-8 sections using the shapes setting in Napari. Then the distance from each detected cell to the pyramidal layer boundary was calculated using a custom Python script based on Chen et al. (2025). To define the pyramidal layer boundaries, reference lines were manually drawn along the SP/SR border in every 5-8 sections. For sections lacking manually drawn reference lines during analysis, cell distances were calculated using the closest section containing a strpyr line. Cell layer classification was defined and based on relative distance from border: superficial layer (−5-22*µ*m); transition zone (22-30*µ*m); deep layer (30-100*µ*m). Cells located below −5 *µ*m were considered off-target (outside the pyramidal layer) and excluded from layer-specific analyses. Analysis included 7 brains (4 right hemisphere, 3 left hemisphere), yielding a total of 3,106 labeled pre-synaptic cells. Cells >100 *µ*m from the border (N=123) and below −5 *µ*m (N=129) were excluded as not part of the stratum pyramidale. A total of N = 2,854 cells were used to study layer distribution. Statistical analyses tested for randomness of distribution using a chi-square goodness-of-fit test across the three layers and compared inputs from deep and superficial layer using two-tailed binomial testing. Effect size was calculated using Cohen’s h, all tests used p = 0.05 as the significance threshold.

### Optogenetic stimulation and in-vitro recordings

Injections were conducted as described under stereotactic injections. Slice prep followed established protocols (Maier et al., 2009). Following anesthesia and decapitation, brains were removed and sliced at 400 *µ*m to preserve hippocampal-cortical connectivity. Slices were stored in an interface chamber (1-6 h) before transfer to the recording setup (ACSF perfusion: 3-4 ml/min). Recordings were made under IR-DIC using an upright microscope. Extracellular and whole-cell patch-clamp recordings targeted PCs in CA1 and SUB (pipette resistance: 2-5 MΩ, multi-patch experiments recorded from up to 8 cells simultaneously. Characterization protocols injected increasing current at steps of 50 pA every 2.5 s. Membrane potential was depolarised or hyperpolarised to keep some cells at −60 mV. Low-resistance pipettes were filled with intracellular solution, which contained (in mM) 120.0 potassium-gluconate, 2.0 MgSO4, 10.0 KCl, 5.0 EGTA, 14.0 Phosphocreatine, 3.0 Mg-ATP, 0.3 Na2-GTP, 10 HEPES buffer, and 0.2% biocytin and the pH was adjusted to 7.2 using KOH. Osmolality was between 275 −290 mOsml/L. Recordings were conducted using Multi-clamp 700B amplifiers (Molecular Devices). Optogenetic stimulation protocols stimulated cells at 10Hz, 10ms, laser intensity was set to spike at 470nm excitation wavelength using the pE – 2 CoolLED Illumination System https://www.microscopy.uk.com/wp-content/uploads/2016/04/CoolLED-pE-2.pdf To test for monosynaptic vs polysynaptic responses in bursting cells, 1 *µ*M TTX was applied; abolished responses that could be recovered with 100 *µ*M 4-AP confirmed monosynaptic Calb-containing CA1 to SUB connections.

### Barnes Maze task and CALB1+ CA1 PC inhibition

Animals completed 4 training rounds (round 1-4), each round starting every two weeks. We ran overall two paradigms (memory consolidation, memory recall paradigm) using a nested-crossover design. Each animal ran through the control as well as the experimental setting of each paradigm. Within each training round, an animal was trained to learn the location of an escape box during 4 training trials (trials 1-4), on 4 consecutive days (training days 1-4). The trainings trials were followed by tests on day 5 and day 12, during which the escape box was removed to assess short-term and long-term memory, respectively. At the start of each trial (duration: 180 s), the mouse was placed in the start box on the center of the platform, after which it had to find the escape box on its own. If the mouse found the escape box before the end of the trial time, the video was ended immediately and the aversive stimuli (white noise of 80 dzb) turned off. If the animal did not find the escape box or entered the escape box, it was gently led towards it. Each animal was kept in the holding box for approx. 15 min between training trials (1-4). For the test days, the escape box was removed and the test lasted 90 s. Afterwards, the mice were collected and returned to the holding box. CNO-induced inhibition of the CALB1+ CA1 PCs was completed in two phases: the first two BM task rounds (rounds 1-2) were designed to study memory recall, by applying CNO during testing, but not during training phase. Experimental and control animals therefore experienced the same conditions during the training phase. The second two BM task rounds (rounds 3-4) were designed to test memory storage, by blocking these cells during the training trials, when the animals were learning a new escape box location. CNO (5mg/kg) and saline (SAL) solution applications took place a minimum of 20 min before the first training trial. CNO was prepared the day before the next trial and kept at 4°C in a non-transparent eppendorf tube to keep the experimenter blinded to the exp/ctrl condition.

### Behavioral Video Analysis

Behavioral videos from 11 CALB1-Cre mice (total N = 784 videos) performing the BM task were analyzed using a custom-built pipeline managed by Snakemake (Mölder et al., 2021), ensuring reproducibility and scalability. The pipeline integrated video pre-processing, deep learning-based tracking, and automated feature extraction to generate the behavioral metrics.

### Video Pre-processing and Arena Definition

Raw video recordings were first processed to correct for recording artifacts. The recordings contained duplicate frames which were removed and temporally corrected for. To exclude artifacts introduced by the presence of the experimenter during initial frames, we implemented an automated procedure to identify and remove frames that exhibit abrupt positional jumps, and then interpolated between the last and the next frame. To define the arena geometry, a mean frame was computed for each video by averaging frames across the recording, excluding the initial period to remove experimenter presence. The locations of the 20 Barnes Maze holes were automatically identified on this mean frame of each video using the `SimpleBlobDetector` from the OpenCV library filtered by area, circularity, and convexity to ensure robust detection.

### Mouse Tracking and Feature Extraction

Mouse body positions were tracked using SLEAP (Pereira et al., 2022). The trajectory of the nose was used as the primary coordinate for behavioral quantification. Tracking data was post-processed to interpolate missing frames and remove artifactual positional jumps. Behavioral metrics were computed based on the tracked nose coordinates and the detected hole locations. A “nose poke” was defined as the nose keypoint entering a 20-pixel radius around the center of a detected hole. Total path length was calculated as the cumulative Euclidean distance of the nose trajectory. To account for camera distortion, an elliptical transformation was applied when converting from pixel to centimeter space. Trial latency was calculated as the time from the start of the trial until the animal entered the escape box. To assess search efficiency versus task completion, latencies were further split into “primary latency” (time to first correct nose poke) and “secondary latency” (time from first correct nose poke to escape box entry).

### Search Strategy Classification

Search strategies were classified for each trial using a hybrid automated-manual approach. First, an automated algorithm analyzed the sequence of nose pokes and center visits to assign a preliminary strategy score based on the classification system described by Rodríguez Peris et al. (2024). Strategies were scored on a scale from 1 (random search) to 11 (direct path), distinguishing between random, serial, and spatial strategies. The automated classifications were manually verified and corrected by a blinded experimenter.

### Spatial Precision Analysis using Von Mises Distribution

To quantify the spatial precision and accuracy of the search behavior, we analyzed the distribution of the first nose poke location relative to the target hole using a von Mises distribution, the circular equivalent of a normal distribution. For each condition and day, the angular positions of the first nose pokes were pooled, and a von Mises distribution was fitted to extract two key parameters: *µ*, representing the mean direction or accuracy of the search, and *κ* representing the concentration or precision of the search. A *µ* close to 0 indicates a search directed towards the target, while a higher *κ* indicates a more precise search pattern with less angular dispersion.

### Statistical Analysis

Statistical comparisons were performed to assess learning and memory performance. Within-group temporal changes (e.g., learning from Day 1 to Day 4, or memory decay from Day 5 to Day 12) were evaluated using the Wilcoxon signed-rank test. Differences in behavioral changes between groups (CNO vs. SAL) were assessed using the Mann-Whitney U test on the calculated change scores. For overall trends across multiple training days, we used Friedman testing. We used permutation testing to assess differences in *µ* between treatment groups. We performed permutation tests on the animal-level parameter estimates (N = 10-11 animals per group). Permutation testing was not done for *κ*, as this is a variability value. For each animal, a single von Mises fit was calculated for the relevant day or condition, yielding one *µ* value per animal. The difference in group means was then compared against a null distribution generated by randomly permuting the group labels 10.000 times.

## Results

### Heterogeneous innervation of both SUB PC types by CALB1+ CA1 PCs

To study whether the laminar segregation of the CA1 extends to the subicular output stage, we combined electrophysiology and monosynaptic rabies tracing to investigate the connectivity between CALB1+ CA1 PCs and SUB PC subtype.

Using Cre-dependent viral constructs targeted to CALB1+ CA1 PCs in CALB-Cre mice, and VGlut2-Cre mice to label the bursting SUB PCs, allowed the mapping of CA1 inputs to distinct SUB cell types and to determine whether CALB1+ CA1 PCs preferentially connect to bursting or regular-firing SUB PCs.

#### Viral construct expression limited to CALB1+ layer of the CA1

As Cavalieri et al. (2021) was able to show that CALB1+ cells are predominantly, but not exclusively, found in the superficial CA1 sublayer, we aimed to determine the specificity of viral expression in CA1 superficial pyramidal cells (CA1 sPCs), and to assess potential off-target expression in the CALB1-CA1 PCs of the deep layer (CA1 dPCs) (Cavalieri et al., 2021). We ran control experiments using a CALB1-CrexAi9 mouse, which globally expressed TdTomato (TdTom) compared to injections and an injection where GFP was only expressed in the soma of Cre-containing cells injected into the dorsal CA1 of the CALB1-Cre mouse-line (Supplementary Fig: 6). We observed expression of mCherry in both superficial and deep layers of the CA1 (Supplementary Fig: 6). This is consistent with the development of the CALB1+ CA1 PCs, with sPCs emerging later than the deep layers, but CA1 dPCs gradually losing CALB1 as the animal matures (Cavalieri et al., 2021). This shows the advantage of targeted viral injections over constitutive reporter lines for selectively accessing CALB1+ CA1 PCs in adult animals. For improved visualization, we tested and optimized existing CALB1 staining protocols (Supplementary Fig: 7).

#### Preferential innervation of VGlut2-containing bursting SUB PCs by CALB1+ CA1 PCs

Using a VGlut2-Cre mouse line which allows the specific targeting of bursting PCs within the SUB, and monosynaptic rabies tracing, we selectively labeled bursting SUB PCs to identify their presynaptic inputs, focusing specifically on the CA1 (Fig: 1A).

**Fig. 1.**
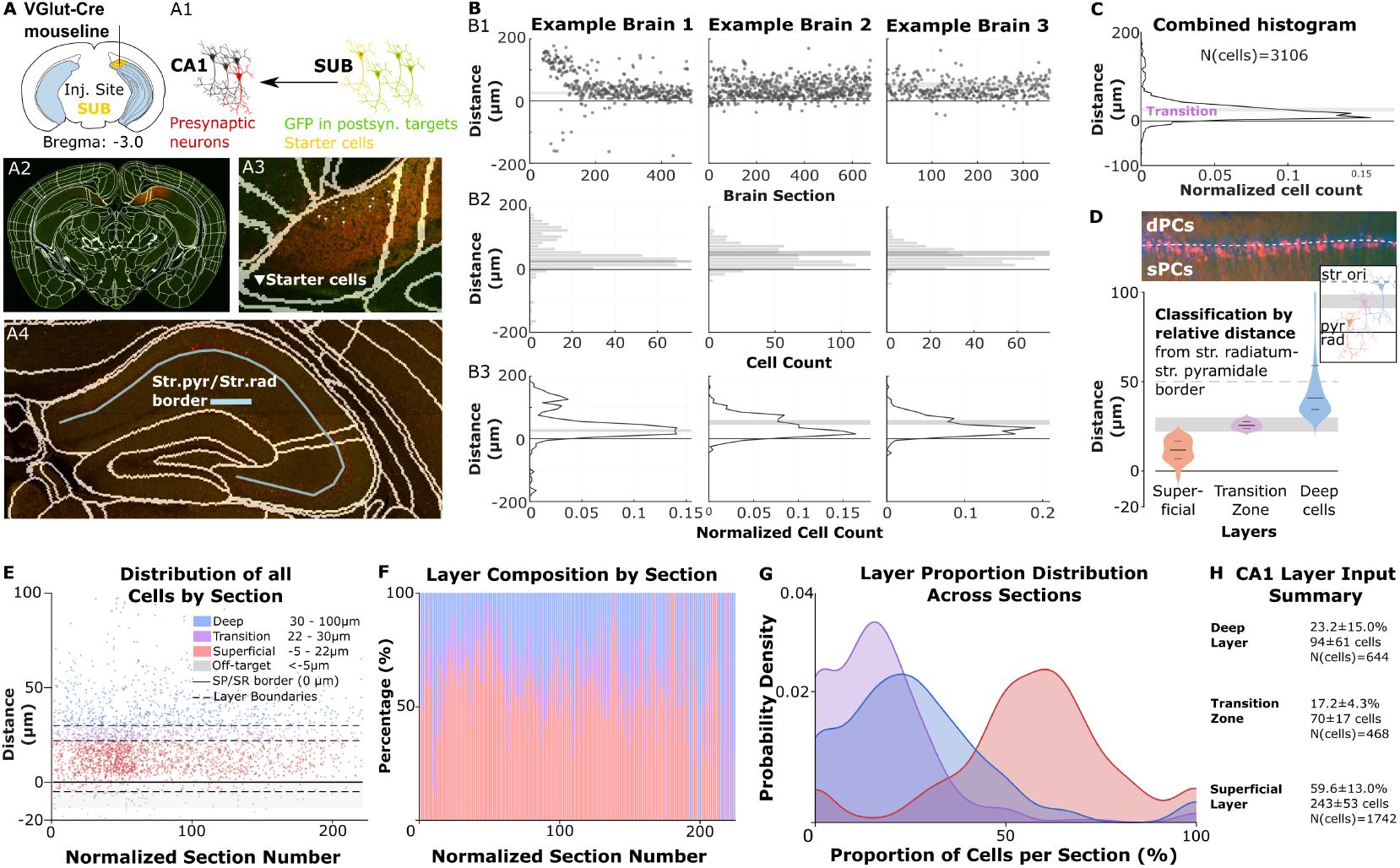
Superficial CA1 PCs preferentially project to VGlut2-bursting SUB PCs. (A) Monosynaptic rabies tracing schematic and representative coronal sections. (A1) Schematic of rabies tracing approach: Injection of AAV-starter and rabies virus into SUB of VGlut2-Cre mice to label starter cells (co-expression of GFP and mCherry, yellow) to see presynaptic input partners in CA1 (red, mCherry). (A2) Whole brain coronal section registered to the Allen Brain Atlas, yellow injection site targets SUB. White lines represent registered atlas boundaries of subareas. (A3) SUB showing starter cells (co-expression of GFP and mCherry, yellow) marked by white arrowheads. (A4) Representative coronal section of the dorsal hippocampus showing mCherry-labeled CA1 PCs. Light blue line indicates manually drawn stratum pyramidale/stratum radiatum (SP/SR) border. (B) Representative cell distribution data from three example brains: (B1) Individual cell positions (raw data) plotted relative to the SP/SR border across normalized sections for each example brain. (B2) Cell count distribution along SP/SR border. (B3) Normalized cell count distribution along the SP/SR border for each example brain. (C) Combined histogram of radial distance from SP/SR border across all datasets (n=7 mice, 2,854 analyzed cells). (D) Example image of mCherry CA1 cells in dorsal hippocampus with layer classification and violin plots. Representative image of mCherry-labeled CA1 PCs in the dorsal HIP, including a line drawn between superficial (sPCs) and deep (dPCs) pyramidal cells based on relative distance from the SP/SR border. Schematic of classification system: superficial layer (red, −5 to +22*µ*m), transition zone (purple, 22 −30 *µ*m), and deep layer (blue, +30 to +100 *µ*m). Violin plots showing layer-specific distribution of cells across all datasets. (E) Distribution of all analyzed cells along normalized sections. Scatter plot of all cells across normalized brain section numbers (superficial-red: −5 to +22*µ*m; transition-purple: 22-30*µ*m; deep-blue: 30 −100*µ*m). Dashed lines indicate layer boundaries. (F) Section composition histogram of superficial, transition, and deep positioned cells across normalized brain sections, displaying percentage composition of each layer across all normalized sections. (G) Probability density curves showing the distribution of cell type proportions per section for superficial (red), transition (purple), and deep (blue) layers. Statistical analysis confirmed a significant bias toward the superficial layer (Chi-square test: *χ*^2^=1001.98, df=2, p<0.001). A direct comparison of superficial vs. deep layers (excluding the transition zone) showed 73.0% superficial and 27.0% deep layer cells (binomial test, p<0.001, n=2,386 cells). (H) Summary of absolute cell counts and pooled percentages for each layer: superficial (1,742 cells, 61.0%), transition (468 cells, 16.4%), and deep (644 cells, 22.6%), out of a total of 2,854 analyzed cells.

This approach allowed us to assess whether the CA1 sPCs preferentially innervate the bursting SUB PC population, or whether these cells receive heterogeneous inputs from both deep and superficial CA1 PCs.

Injections of the starter and rabies virus into the VGlut2-Cre expressing bursting SUB PCs resulted in robust expression of mCherry in CA1, with the majority of the labeled cells located in the CA1 superficial sublayer, although a proportion of labeled cells could also be seen in the deep layer (Fig: 1A1, A2, A3). To quantify the spatial distribution of rabies-traced CA1 PCs, mCherry-expressing somata and their positions relative to the strpyr border were analyzed (Fig: 1A4; see methods). Of a total of 3106 labeled pre-synaptic cells within the CA1 of the hippocampus, 2854 CA1 PCs were analyzed.

Analysis classified 59.6 ± 13.0% (N=1742) as belonging to the superficial, 17.2 ± 4.3% (N=468) to the transition, and 23.2 ± 15.0% (N=644) to the deep layer (N=7 mice) (Supplementary Fig. 7). 4.3% (129 cells) were located below −5 *µ*m and excluded, as they were not based within the strpyr layer. To avoid misclassifications, a transition zone was defined as 22-30*µ*m to account for the gradual transition between superficial and deep layers, as most publications have primarily drawn a boundary between these two layers based on their relative distance from the strpyr border (Sharif et al., 2021; Valero et al., 2015; Geiller et al., 2017; Hainmueller and Bartos, 2018) (Fig: 1C, E). The chi-square goodness-of-fit test showed that labeled cells were not randomly distributed along the width of the strpyr layer (*χ*^2^ = 1001.98, p < 0.001). A direct comparison of labeled PCs within the superficial versus deep layer (excluding 468 cells within transition zone) showed that sPCs were significantly more likely to project towards the VGlut2-SUB PCs (1742/2386 cells, 73.1%) compared to deep cells (644/2386 cells, 26.9%; binomial test, p < 0.001).

This superficial bias was consistent across all 7 animals analyzed. This shows that though both layers project to the VGlut2-SUB PCs, there is a preference with a sPC:dPC ratio of 2.7:1 (Cohen’s h=0.96). Removing the transition zone and using a single boundary line at 25*µ*m (midpoint of the 50-55*µ*m pyramidal soma layer (Sharif et al., 2021; Cavalieri et al., 2021) still showed a significant bias of inputs from the superficial layer (Fig: 1E, G, H): superficial inputs made up 67.2%, and deep layer 32.8%, a ratio of 2.05:1 (binomial test, p < 0.001, Cohen’s h=0.7). This shows that although the bursting SUB PCs are innervated by both layers of the CA1, the preference of innervation by the superficial CA1 layer remains robust.

#### CALB1+ CA1 PCs induce spiking in both bursting and regular-firing SUB PCs and more reliably in bursters

To study the physiological importance of the connection between CA1 sublayers to SUB PC subtypes, we conducted single-cell patching experiments. By patching pyramidal cells in the SUB in slices expressing Channelrhodopsin (ChR2) in the terminals of CALB1+ CA1 PCs in the dorsal hippocampus (Fig: 2A1, A2) and optogentically stimulating (10 Hz, 10 ms) the fibers of the ChR2-expressing cells in the SUB, we were able to assess whether activation of CALB1+ CA1 PCs was able to induce spiking in the SUB PCs. GFP was expressed in the terminals (Fig: 2A3, A4, A5). Prior to optogenetic stimulation, we ran characterization protocols to assess the SUB cell type. Cells were patched in the medial SUB to study both regular and burst firing cells, as patching from only distal or proximal SUB would have skewed which population type would be most abundant -as shown by (Cembrowski et al., 2018; Wozny et al., 2018), the regular-firing PCs can be found mainly on the proximal, while burst-firing PCs are mainly found in the distal end of the SUB. Additional multi-patching experiments allowed us to study whether this same pattern could be observed simultaneously within the same slice, as well as assess whether these connections were mono-or polysynaptic (Fig: 2B). To do this, we conducted whole-cell patch-clamping on PCs within the medial SUB. Across all patching experiments, we recorded a total of 29 SUB PCs cells (N=15 bursters, N=14 regular-firing). After optogenetic activation of the ChR2-expressing terminals of CALB1+ CA1 PCs, 15 SUB PCs in total spiked in response to light stimulation, of which 12 were burst-firing PCs (80% of all recorded bursters) while only 3 were regular-firing cells (21% of all recorded regular-firing cells) (Fig: 2C).

**Fig. 2.**
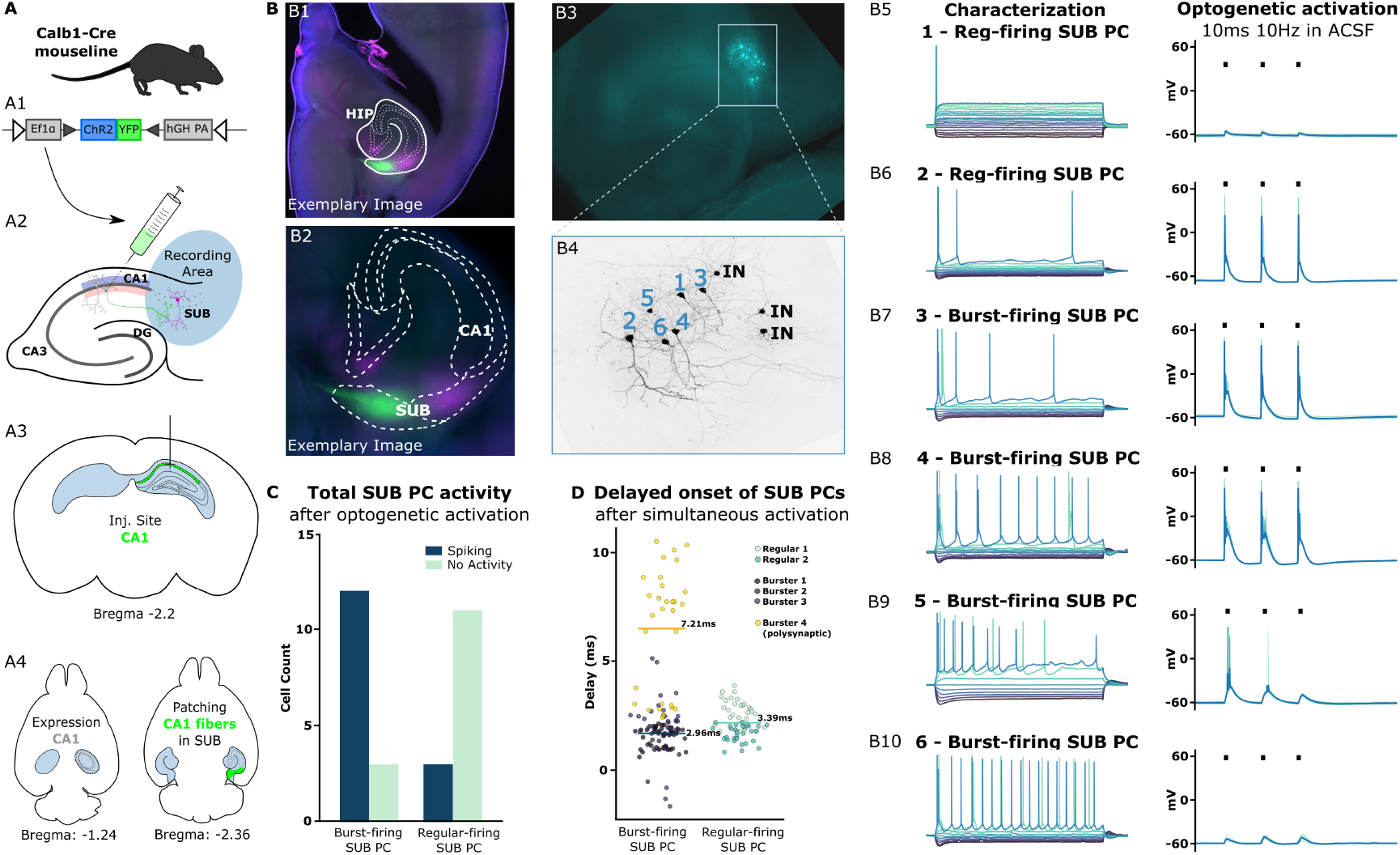
CALB1+ CA1 PCs induce spiking more reliably in burst-than regular-firing SUB PCs. (A) Experimental approach. (A1) Viral construct (pAAV-EF1a-Switch-mRubyNLS-hChR2(H134R)-EYFP-NO-WPRE) injected into CALB1-Cre mice to express ChR2-GFP in CALB1+ CA1 PCs. (A2) Schematic of hippocampus (coronal view) detailing sublayers of CA1, location of ChR2-expression and and patching location in dorsal SUB with optogenetic stimulation. (A3) Injection site at bregma −2.2 mm. (A4) Transverse sections showing injection site within CA1 (bregma −1.24 mm) and GFP-linked ChR2-expressing in CA1 terminals ending in the SUB (bregma −2.36 mm, green). (B) Exemplary multipatch experiment. (B1) Whole hippocampal slice showing GFP expression in CA1 fibers within the SUB. (B2) Close-up of CA1 and SUB regions. (B3) Post-recording image of Alexa-stained patched cells in medial SUB. (B4) Close-up of patched PCs: 4 burst-firing, 2 regular-firing, 3 fast-spiking interneurons (IN). (B5-B10) Recorded SUB PCs (characterization trace (left), followed by light-evoked responses (10 ms, 10 Hz) in ACSF, regular-firing (reg-firing) and burst-firing SUB PCs. Exclusion of INs. (C) Summary of all recorded SUB PCs across all experiments an their response to light stimulation in ACSF. Spiking PCs (dark blue): 12/15 bursters (80%), 3/14 regular-firing (21%). No activity (light green: 3/15 bursters, 11/14 regular-firing. (D) Onset delay analysis across 10 sweeps for all PCs recorded in multipatch experiment shown in (B). Onset delay: bursters (dark blue) mean 2.96 ms; regular-firing (light green) mean 3.39 ms; one burster (yellow) shows prolonged latency and could not be rescued with 4AP, mean 7.21 ms. Mean onset delays indicate predominantly monosynaptic connectivity.

#### Connection of CALB1+ CA1 PCs to SUB PCs are largely monosynaptic

To test whether these inputs were mono-or polysynaptic, we tested the response of a proportion of cells to light stimulation after application TTX, to suppress the activation and spiking of these cells, and then added 4AP to rescue monosynaptic connections from the CALB1+ CA1 PCs to the SUB PCs (Fig: 3B5, B6). Across 3 slices, we applied these conditions to 6 burst, and 3 regular firing cells overall. In one specific multipatching experiment, we simultaneously recorded from 6 SUB PCs within the same slice (N=4 bursters, N=2 regular-firing). One burster and one regular-firing PC were lost between the initial ACSF recordings and the TTX deactivation and rescue protocols. Combined with the single-cell recordings, we were able to observe that the activation profiles of 4/6 bursters and the remaining regular-firing PC could be rescued, indicating monosynaptic connec-tions. The burster, which could not be rescued after 4AP application was classified as a polysynaptic connection (Supplementary Fig: 8). These findings are supported by analysis of the delayed onset after light stimulation of the cells recorded in ACSF baseline experiments. The subpopulation of bursters that could be rescued by use of 4AP showed rapid onset delays of 2.62 ± 0.61 ms (N=3 bursters), and the regular-firing PCs showed an onset delays of 3.39 ± 0.64 ms (N=2 regular) (Fig: 2D). Meanwhile, the burster that could not be rescued by application of 4AP also displayed a longer onset delay of 7.21 ± 2.42 ms. Recordings were conducted in the presence of Gabazine to ensure exclusive activation of excitatory pyramidal cells.

**Fig. 3.**
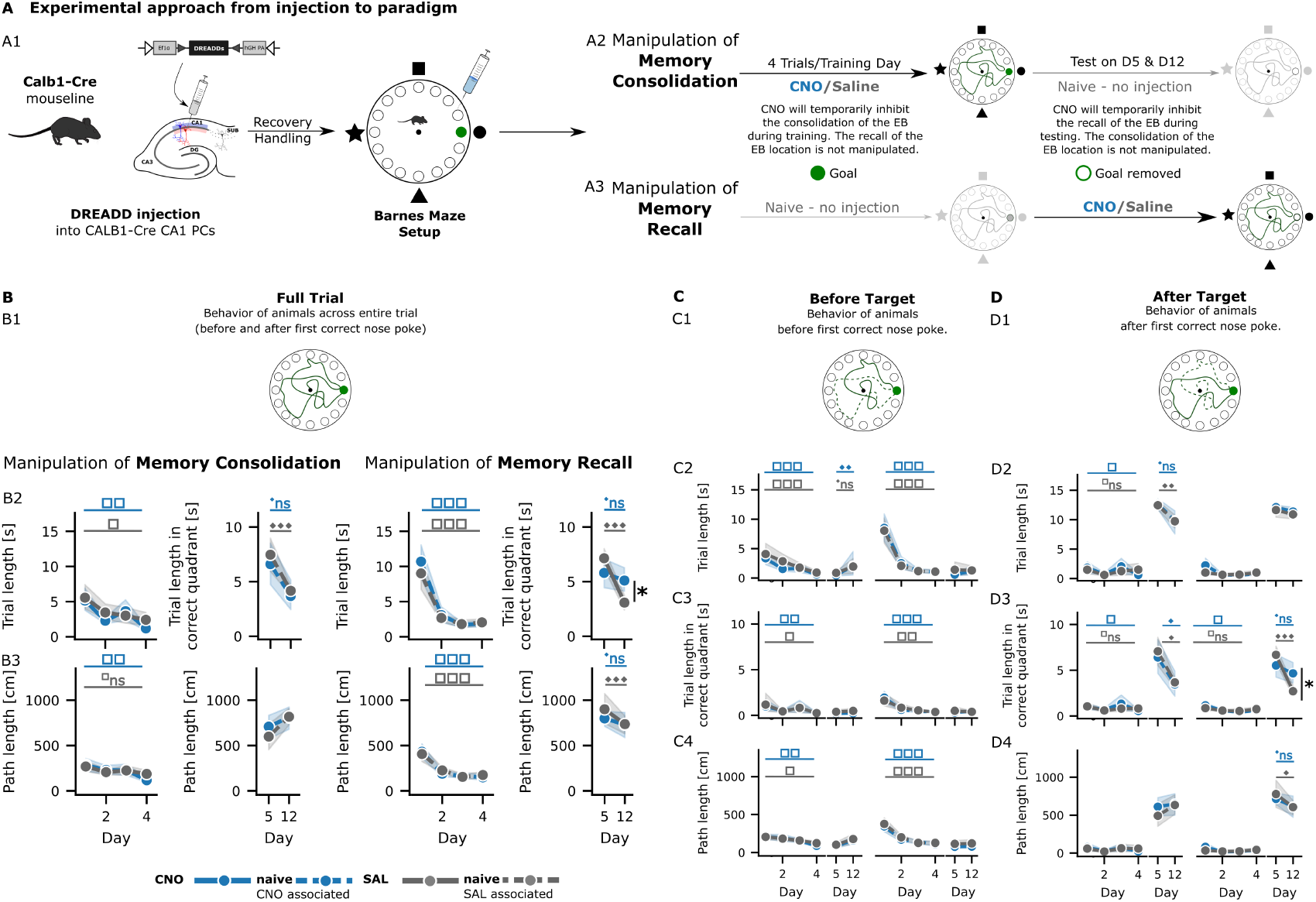
Inhibition of CALB1+ CA1 PCs does not impair general learning or locomotor abilities and lowers cognitive flexibility. (A) Overview of experimental approach. (A1) Handled CALB1-Cre animals received Cre-dependent DREADD injections into dorsal CA1. After recovery, animals were exposed to the Barnes Maze (BM) setup. During training days (Days 1-4), animals escaped via a target escape box (EB, green). The EB was removed during testing days (Day 5 STM, Day 12 LTM), with no BM exposure between testing sessions. CNO was injected to inhibit CALB1+ CA1 PCs, saline (SAL) for control. (A2) Memory consolidation paradigm: CNO/SAL given before each training session to manipulate memory consolidation. Memory recall paradigm: CNO/SAL given before each testing session to manipulate memory recall. EB location changed between paradigms. (B) Full trial BM analysis. (B1) Schematic of full trial, animal approaches correct hole, and then continues to explore before entering EB. Data show complete trial duration from start until EB entry. CNO (blue), SAL (gray), with naive controls shown as dashed lines. Error bars represent 90% confidence intervals. (B1) Full trial length during testing phase. Full trial length in correct quadrant shown in testing phase. Memory consolidation paradigm (panels 1-2): Both groups show equivalent learning during training. Memory recall paradigm (panels 3-4): Groups differ significantly on Day 12 (ANOVA p < 0.05), with CNO animals showing reduced cognitive flexibility to look for new hole. (Friedman p < 0.05 for all training phases). (B3) Full trial path length mirrors latency findings, confirming learning across training phases. (C) Before-target analysis. Behavior measured from trial start until first correct nose poke. (C1) Schematic showing behavior before first correct nose poke. (C2) Trial length before first correct nose poke shows significant learning effects (Friedman p < 0.05 for all training phases). (C3) Trial length in correct quadrant before first correct nose poke. (C4) Path length before first correct nose poke demonstrates spatial learning efficiency across both groups (Friedman p < 0.05 for all training phases). (D) After-target analysis. Behavior measured from first correct nose poke until EB entry. (D1) Schematic showing behavior after first correct nose poke. (D2) Trial length after first correct nose poke. (D3) Trial length in correct quadrant after first correct nose poke reveals impairment of cognitive flexibility in CNO group during memory recall testing (significant group difference on Day 12, ANOVA p < 0.05). (D4) Path length after first correct nose poke shows path length of CNO-group remains the same from STM to LTM test. CNO in blue, SAL in gray, dashed lines-non-injected group; line-mean, shaded error line −STD. Statistical annotations: ANOVA between conditions: * p < 0.05, p < 0.01, * p < 0.001. Friedman across days within group: □ p < 0.05, □□ p < 0.01, □□□ p < 0.001. Wilcoxon between days within group: ♦ p < 0.05, ♦♦ p < 0.01, ♦♦♦ p < 0.001. Whitney U between groups: × p < 0.05. Not significant is shown as n.s. or not shown if not significant for all respective comparisons.

Characterization and subsequent light stimulation of fastspiking interneurons (N=9 interneurons) confirmed that these were not involved in the e.g. polysynaptic activation of SUB PCs. Control experiments, which used a virus expressing ChR2 in all CA1 PCs led to spiking in all recorded SUB PCs (N=2 bursters; N=4 regular-firing).

Both the rescue experiments and onset delay analysis support the notion that connections from CALB1+ CA1 PCs to both SUB PC populations are largely monosynaptic, though polysynaptic connections cannot be excluded. This aligns with our retrograde rabies tracing findings: the preferential anatomical connectivity from the superficial, largely CALB1+ CA1 PC layer to burst-firing SUB PCs translates functionally into more reliable spike induction in bursters than in regular-firing cells. Both approaches indicate a preferential, though not exclusive, connection between these two PC subpopulations of the CA1 and SUB.

### Effect of inhibition of CALB1+ CA1 PCs in Barnes Maze Task

Given that we know that the inhibition of the distal SUB, which is known to primarily harbor bursting SUB PCs, disrupts the consolidation of a spatial memory task such as the MWM, we next wanted to study the implications of inhibiting the CA1 subpopulation preferentially innervating them: the CALB1+ CA1 pyramidal cells found in the superficial layer on the radial axis (Cembrowski et al., 2018). We inhibited the CALB1+ CA1 PCs in either the training or testing phase of a memory consolidation and recall paradigm, respectively, to test their functional relevance. The memory consolidation paradigm used CNO-inhibition at the start of each day of the training phase, potentially impacting memory consolidation. In the recall paradigm, animals consolidated the location of the goal location naively (non-injected), and only received CNO-inhibition on testing days (Fig: 3A2, A3). We first focused on the classical analysis of the Barnes Maze by analysing the changes of time and distance parameters as the animals navigated the platform to evaluate their learning and memory.

#### CALB1 CA1 PC inhibition affects latency and path length, lowers cognitive flexibility

Crucially, in the memory consolidation paradigm, a clear decline of total trial latency could be observed in both groups across training days (D1(SAL) 5.6 ± 6.6 to D4(SAL) 2.4 ± 3.8 s, Friedman p = 0.048, N=44; D1(CNO) 5.2 ± 6.5 to D4(CNO) 1.2 ± 1.3 s, Friedman p = 0.004, N=44), indicating learning (Fig: 3B2). Primary latency analysis also showed that both groups reduced the time required to learn the correct goal location in the training phase (primary latency: D1(SAL) 4.1 ± 6.3 to D4(SAL) 0.9 ± 1.0 s, Friedman p = 0.01, N=44; D1(CNO) 3.3 ± 4.8 to D4(CNO) 0.6 ± 0.8 s, Friedman p = 0.001, N=44) (Fig: 3C2). Experimental and control groups did not differ in their learning on each respective training day (D1-D4 ANOVA p > 0.05). Meanwhile, differences could be observed when studying secondary latency, the time between the first correct nose poke and entering the escape box: this time remained the same for the SAL-injected group during the training phase (secondary latency: D1(SAL) 1.5 ± 3.0 to D4(SAL) 1.5 ± 3.7 s, Friedman p > 0.05, N=44), while CNO-injected animals continued to reduce their secondary latency (D1(CNO) 1.8 ± 4.0 s to D4(CNO) 0.6 ± 1.0 s, Friedman p = 0.033, N=44) (Fig: 3D2). This could also be shown using a Wilcoxon test, which revealed that CNO animals reduced their secondary latency (CNO, Wilcoxon p = 0.027) during the training phase from Day 1 to Day 4, while control animals showed no significant change (SAL, Wilcoxon p = 0.898), indicating that CNO animals became more efficient at completing the task after locating the target during the training phase. Changes could be observed in the testing phase of the consolidation paradigm: Both groups reduced the time spent in the correct quadrant from the short-term memory test on D5 (STM) to the long-term memory test on D12 (LTM). Wilcoxon tests reveal both groups showed significant intra-group changes: SAL-injected animals (D5(SAL) 7.5 ± 3.1 (N=10), to D12(SAL) 4.2 ± 1.0 (N=11) s; Wilcoxon p = 0.014) reduced the time spent in the correct quadrant by 44.1%. This was the same for animals injected with CNO during the training, but not the testing phase, as they also reduced the time spent in the correct quadrant by 44.1% D5(CNO) 6.6 ± 3.7 (N=10) to D12(CNO) 3.7 ± 2.6 (N=9) s, Wilcoxon p = 0.02).

In the memory recall paradigm, learning was also indicated by the same reduction in total trial latency in both groups across training days (D1(SAL) 9.0 ± 9.2 to D4(SAL) 2.0 ± 2.0 s, Friedman p < 0.001, N=44; D1(CNO) 10.7 ± 9.4 to D4(CNO) 1.8 ± 1.9 s, Friedman p < 0.001, N=44) (Fig: 3B2). Primary latency also showed that both groups were able to reliably reduce the time required to find the correct goal location in the training phase. Neither group was injected during the training phase, and both groups behaved the same on each respective training day (D1 to D4, ANOVA p > 0.05) (Fig: 3C2). Unlike in the memory consolidation paradigm, differences were not observed between groups in regard to secondary latency: both SAL-injected animals (D1(SAL) 1.0 ± 1.3 to D4(SAL) 1.0 ± 1.4 s, Friedman p = 0.427, N=44), and CNO-injected animals (D1(CNO) 2.2 ± 4.6 s to D4(CNO) 0.8 ± 1.3 s, Friedman p = 0.071, N=44) did not significantly alter their behavior after the first nose poke (Fig: 3D2). Meanwhile, changes could be observed in the testing phase of the recall paradigm, which is when CNO/SAL was injected for the first time: Though both groups reduced the time spent in the correct quadrant from STM to LTM, the two groups differed significantly from one another in the LTM (ANOVA p = 0.021). Component analysis showed that control animals displayed a marked drop of 56.9% in time spent in correct quadrant between testing days (D5(SAL) 7.1 ± 1.9 (N=11), to D12(SAL) 3.1 ± 1.4 (N=11) s; Wilcoxon p < 0.001). CNO-injected animals on the other hand remained in the correct quadrant, reducing the time spent here only by 12% between STM and LTM (D5(CNO) 5.8 ± 2.8 (N=11) to D12(CNO) 5.1 ± 2.3 (N=11) s, Wilcoxon p > 0.05) (Fig: 3D3). This indicates that while there was a loss in accuracy from STM to LTM for both groups, CNO-injected animals maintained their focus on the correct location for longer, while SAL-injected animals were faster to pivot to renewed exploratory behavior.

Overall, path length analysis (Fig: 3B3, B4, D4) supports our latency findings. Both groups display similar increased efficiency and reduced path length in the training phase of both consolidation and recall paradigm. For example, control animals decreased by 29.9% (D1(SAL) 266.5 ± 37.4 to D4(SAL) 186.9 ± 30.6 cm, Friedman p = 0.167); CNO-injected animals decreased by 56.8% (D1(CNO) 284.0 ± 37.3 to D4(CNO) 114.9 ± 19.3 cm, Friedman p = 0.003) in the training phase of the consolidation paradigm (Fig: 3B3). Just like for the total path length during test phases in both paradigms, analysis showed no differences when looking at the behavior of both groups before navigating to the escape box (Mann-Whitney U, all p > 0.05), indicating that inhibition of only CALB1+ CA1 PCs does not impact locomotor control and general motor activity, as opposed to a global CALB1-KO (Airaksinen et al., 1997; Farré-Castany et al., 2007).

In summary these findings indicate that when CNO is administered in the training phase of the memory consolidation paradigm, both groups largely behave the same in training and testing phases. Meanwhile, CNO-administration during the testing phase of the recall paradigm affects cognitive flexibility.

### Cognitive performance of animals is influenced by inhibition of CALB1+ CA1 PCs

While latency and path length are useful to assess the performance of an animal in the Barnes Maze task, these parameters do not capture cognitive performance. To further understand the effect of CALB1+ inhibition on spatial learning, we examined changes in search strategy and the effect on learning precision and perseverance in what had been learned.

#### No impairment in Barnes Maze search strategy development and execution

Search strategy was quantified by adapting a novel search strategy classification system (Rodríguez Peris et al., 2024) and implementing an automated scoring approach across all videos. This scoring system acts as a sensitive tool to study contextual and procedural strategies to reach the correct goal location on a range from 1 (random, non-sequential hole checking) to 11 (direct trajectory to target without checking other holes). Higher scores (red, orange, yellow, green) indicate contextual search strategies, while lower scores reflect more procedural or exploratory behavior (dark blue, purple) (Fig: 4A). The search strategy scores captures the behavior during the primary search phase, thereby describing the behavior before the first correct nose poke.

In the training phase of the memory consolidation paradigm, SAL-injected animals did not significantly alter their strategy scores (D1(SAL) 6.26 ± 0.47 to D4(SAL) 7.45 ± 0.42, N=44, Friedman p > 0.05), while CNO-injected animals showed significant improvement (D1(CNO) 6.2 ± 0.41 to D4(CNO) 8.08 ± 0.43, N=40-44, Friedman p = 0.039). It must be noted that no significant differences could be observed between both groups on individual training days (D1-D4 ANOVA p > 0.05) (Fig: 4B1). During the testing phase, primary search strategies did not significantly differ between STM and LTM for either group, as scores remained high for control animals (D5(SAL) 7.4 ± 1.0 to D12(SAL) 7.2 ± 0.8, N=10-11, Wilcoxon p > 0.05), and for CNO-injected animals (D5(CNO) 8.2 ± 0.9 to D12(CNO) 7.7 ± 1.1, N = 9, Wilcoxon p > 0.05), indicating that both groups remained committed to corrections (score 7) as their average search strategy.

In the memory recall paradigm, both groups showed significant strategy improvement during the training phase: both SAL-injected animals (D1(SAL) 4.5 ± 0.5 to D4(SAL) 7.7 ± 0.5, N=44, Friedman p = 0.013) and CNO-injected animals (D1(CNO) 4.7 ± 0.5 to D4(CNO) 8.6 ± 0.4, N=44, Friedman p = 0.001) adapted their search strategies from serial (score 4) to corrections (score 7) across all four training days (Fig: 4B2). Yet again, both groups remained committed to their primary search strategies, as both groups did not alter their behavior significantly between STM and LTM (Wilcoxon p > 0.05).

**Fig. 4.**
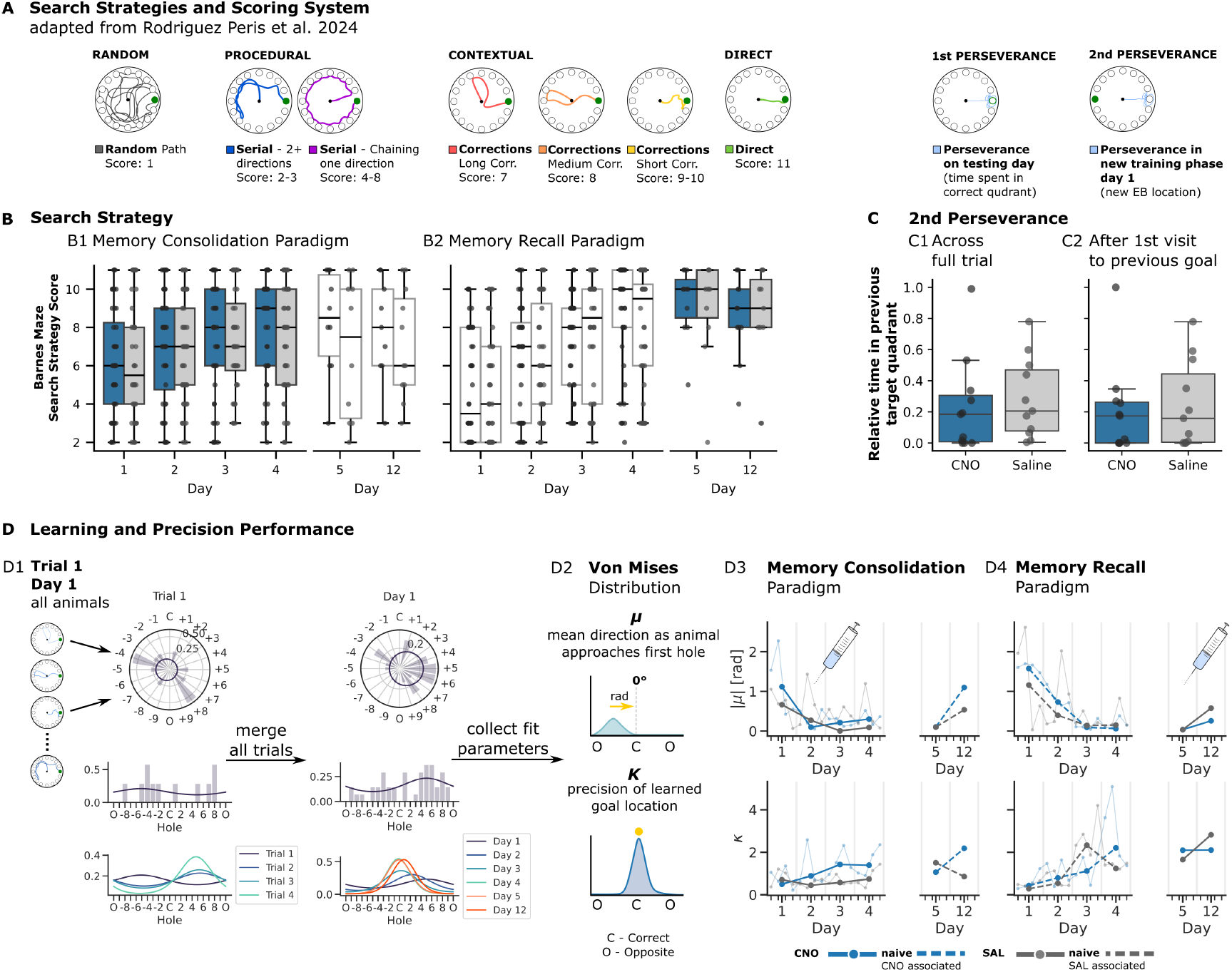
Search Strategy Classification and Cognitive Performance Analysis using Von Mises Distribution. (A) Search strategy classification system using automated implementation of Rodriguez-Peris et al. (2024) scoring system. Representative tracks show: random strategy (score 1, dark grey), serial strategies with direction changes (scores 2-3, dark blue) or unidirectional chaining (scores 4-8, purple), and contextual strategies including long corrections (score 7, red), medium corrections (score 8, orange), short corrections (scores 9-10, yellow), and direct search (score 11, green). Perseverance shown as 1st level perseverance (time spent in correct quadrant on test days when the EB (green) is removed) and 2nd level perseverance (time spent in quadrant of EB location of the previous round during training phase of the new round, EB has been relocated (green circle). (B) Search Strategy classification across both memory consolidation and recall paradigm. (B1) Strategy use in memory consolidation paradigm: groups shift from procedural to contextual strategies during training, both groups use predominantly contextual strategies in STM, and revert to procedural strategies in LTM, no difference between groups (ANOVA p > 0.05). (B2) Strategy use in memory recall paradigm: shift from procedural to contextual strategies during training, both groups use predominantly contextual strategies in STM and LTM. (C) 2nd level perseverance for both groups, no difference between groups (ANOVA p > 0.05). (C1) Relative time in previous target quadrant across full trial, no difference between groups (ANOVA p > 0.05). (C2) Relative time in previous target quadrant after first poke of previously correct goal location, no difference between groups (ANOVA p > 0.05). (D) Learning and precision performance captured by von Mises distribution. (D1) von Mises distribution represents analysis of the absolute first nose poke: trials of all animals are pooled to generate von Mises fits for individual trials (1st row: polar representation; 2nd row von Mises fit of single trial, all animals; 3rd row: von Mises fits for all trials, day 1), from which all trials are merged for von Mises distribution for individual day (1st row: polar representation; 2nd row: von Mises fit D1; 3rd row: von Mises fits for all days combined). (D2) Schematic of von Mises parameters: *µ* (mean direction animal heads towards for first hole poked; C = correct hole (0), O = opposite of correct hole), *κ* (mean distribution of first hole poked, describing precision of animal’s learning). (D3) von Mises fit parameters in memory consolidation paradigm, *µ* decreased during training phase, remained low on D5, increased for CNO on D12; *κ* increased during training for CNO more than SAL; CNO showed higher *κ* on D12 than SAL. (D4) Von Mises fit parameters in memory recall paradigm: *µ* decreased in both groups during training phase, remained low on D5, increased more for SAL on D12; *κ* increased for both groups during training, SAL scores higher on D12. Permutation testing revealed no significant differences in *µ* between groups. (CNO (N=11 animals),SAL (N=11 animals), dashed lines: non-injected controls.)

As the Barnes maze provides circular data, it is possible to examine the distribution of holes visited during a trial using a von Mises distribution. This probability density function allows for the analysis of directional data (see methods for more information). The parameters *µ* and *κ* represent the mean and variance in a normal distribution, respectively. In the case of the Barnes maze task, *µ* represents accuracy of the mean search direction, and displays the primary search direction relative in radius to the correct hole. The parameter *κ* focuses on the peak of the distribution, and functions as an indicator of precision. A high *κ* describes highly precise, goal-directed approach behavior. The von Mises fit were captured by pooling all animals across trials and days, and fitting the distribution of their first nose poke relative to the correct hole location (Fig: 4D). This approach allowed us to extract *µ* and *κ* parameters that quantify both the accuracy and precision of spatial memory across training and testing phases (Fig: 4D1, D2).

In the training phase of the memory consolidation paradigm, both groups showed improved accuracy, as *µ* successively centered towards the correct hole from D1 (D1(SAL) 0.66 rad to D4(SAL) 0.09 rad, N=44; D1(CNO) 1.12 rad to D4(CNO) 0.30 rad, N=44). This supports our findings of increasing contextual,increasingly direct strategy use (Fig: 4B1). During the STM test, both groups displayed similar accuracy (D5(SAL) 0.10 rad, N=11; D5(CNO) 0.10 rad, N=11), but during the LTM test, animals treated with CNO during the training phase approached their first hole with less directional accuracy than the SAL group (D12(SAL) 0.54 rad, N=11; D12(CNO) 1.10 rad, N=11) (Fig: 4D3). The *κ* parameter, an indicator of precision in learning, increased for CNO-injected animals during training (D4(SAL) 0.74, N=44; D4(CNO) 1.39, N=44). Here, animals which had previously been treated with CNO also increased their scores between the STM and LTM test (D12(SAL) 0.85, N=11; D12(CNO) 2.19, N=11), indicating that although CNO animals were less accurate in terms of mean search direction, they were more confident in their search behavior.

In the memory recall paradigm, both groups showed similar trajectories during the training phase (D1(SAL) 1.16 rad to D4(SAL) 0.15 rad, N=44; D1(CNO) 1.58 rad to D4(CNO) 0.06 rad, N=44) (Fig: 4D4). During the STM test, both groups also showed similar accuracy (D5(CNO/SAL) 0.04 rad), while SAL-injected animals initiated search in directions further from the correct goal location than CNO animals during the LTM test (D12(SAL) 0.58 rad, N=11; D12(CNO) 0.26 rad, N=11), potentially reflecting our findings in the latency analysis (Fig: 3B2, D3). Meanwhile, the *κ* values for the control animals were higher on the last testing day (D12(SAL) 2.82, N=11; D12(CNO) 2.10, N=11). To assess the statistical significance of these observed differences in *µ* between groups, we performed permutation testing, which revealed no significant differences between groups (all p > 0.05). Permutation testing did reveal a higher accuracy for all injected animals (CNO and SAL) versus the non-injected (naive) animals of the same phase of the other respective paradigm, suggesting that the stress of the injection encourages animals to find the escape box accurately.

## Discussion

The hippocampus has been shown to be functionally specialized along each axis. While CA1 sublayers differ in their physiological properties and computational roles (Danielson et al., 2016; Mizuseki et al., 2011; Esparza et al., 2025; Geiller et al., 2017; Sharif et al., 2021), it remains unknown whether this laminar organization extends to their connections to the SUB, the hippocampal output hub through which processed information reaches cortical networks. This question is relevant given that bursting SUB PCs have been shown to propagate SWRs to the gRSC (Nitzan et al., 2020) and have also been shown to be necessary for spatial memory (Cembrowski et al., 2018). We therefore investigated a potential microcircuit between genetically defined CALB1+ CA1 PCs and SUB PC subtypes using retrograde monosynaptic rabies tracing, slice electrophysiology and patch-clamping, and chemogenetic manipulations in a spatial memory task. Our findings reveal that CALB1+ CA1 PCs in the superficial layer of the radial CA1 axis preferentially connect to bursting SUB PCs and that this pathway seems to support cognitive flexibility, while not necessary for initial learning and memory consolidation.

### Functionally and anatomically relevant microcircuit between CALB1+ CA1 PCs and bursting SUB PCs

Retrograde monosynaptic rabies tracing from VGlut2-expressing bursting SUB PCs revealed that 73% of presynaptic CA1 inputs originated from the superficial layer while only 27% came from the deep layer, yielding a 2.7:1 ratio. These findings remained consistent with a boundary criterion (67.2% superficial vs. 32.8% deep, 2.05:1 ratio, Fig: 1E, H). Electrophysiological experiments functionally supported this anatomical preference: 80% of bursting neurons (12/15) responded with spiking, compared to only 21% of regular-firing cells (3/14). Short onset latencies (2.96 ms for bursters, 3.39 ms for regular-firing cells) and successful rescue following TTX application with 4-AP confirmed predominantly monosynaptic connectivity (Fig: 2). These findings show that the laminar segregation along the radial axis of the CA1 extends to its output pathways, as CALB1+ CA1 PCs form a preferential microcircuit with bursting SUB PCs.

### Potential microcircuit mechanisms of spatial map establishment and transfer to cortical areas

The electro-physiological, genetic, and developmental differences between CA1 sublayers (Soltesz and Losonczy, 2018) are increasingly linked to distinct functional roles (Esparza et al., 2025; Johantges et al., 2025). Based on the preferential connectivity we observed, we propose that CALB1+ CA1 PCs are positioned to be regulators of hippocampal output in a larger microcircuit between interneurons and PC subtypes of the CA1 and SUB (Fig: 5). This is based on recent findings which showed that superficial CA1 PCs, including CALB1+ cells, form reciprocal connections with somatostatin-positive interneurons (SST+ INs), while deep layer CA1 PCs receive stronger inhibition from parvalbumin-positive basket cells (PV+ BCs). This supports an inhibitory feedback loop within the CA1 (Johantges et al., 2025), a feedback mechanism which might enable the temporal gating of signals to the SUB through delayed, dendrite-targeted inhibition. This in turn allows for increased control over CA1 activity, and relay to downstream targets (Johantges et al., 2025). The preferential connectivity from CALB1+ CA1 PCs to bursting SUB PCs, combined with distinct inhibitory circuits for the CA1 PC sublayers could act as a precision control mechanism, regulating the amplitude, timing, and duration of excitation to SUB bursting cells. This is relevant given that burst-firing SUB PCs are essential for the propagation of SWRs, and critical for memory consolidation (Nitzan et al., 2020; Cembrowski et al., 2018). Within the SUB, the unidirectional connectivity from regular-firing to burst-firing SUB PCs could enable the SUB to further integrate inputs before propagating consolidated information to cortical targets. (Böhm et al., 2015). A recent study has also observed a higher participation of the superficial layer of the CA1 in SWR propagation, supporting our proposed microcircuit (Berndt et al., 2023).

**Fig. 5.**
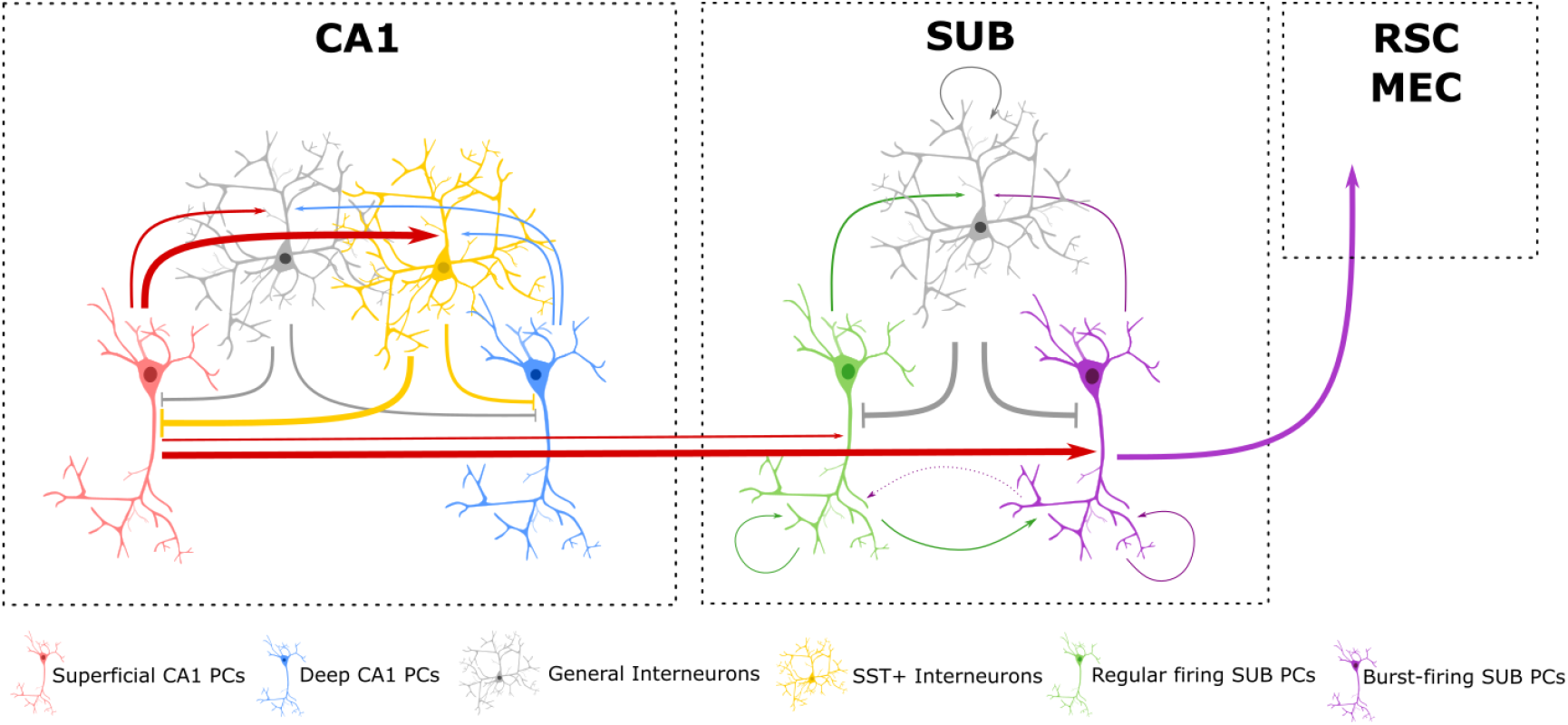
Proposed microcircuit between CA1 sublayers within the CA1, the SUB and Cortical Areas. Superficial CA1 PCs (red) preferentially connect with somatostatin-positive interneurons (SST+ INs, yellow), while the CA1 of the deep layer (blue) receive stronger inhibition from Parvalbumin-positive basket cells (PV+ BC) (Johantges et al., 2025). CALB1+ CA1 PCs preferentially target burst-firing, VGlut2-positive SUB PCs (purple) over regular-firing SUB PCs (green). Within the SUB, burst-firing and regular-firing PCs are involved in local microcircuits with each other and interneurons (gray), and display exclusive unidirectional connectivity from regular-firing SUB PCs to burst SUB PCs, linking functional specialization to local network topology (Böhm et al., 2015). Burst-firing SUB PCs have additionally been shown to project to the granular retrosplenial cortex (gRSC) and other cortical targets (Nitzan et al., 2020). The preferential connection of superficial CA1 PCs to SST+ INs might be able to create an inhibitory feedback loop which could support temporal gating of subicular activation through delayed, dendrite-targeted inhibition (Johantges et al., 2025). In tandem with the preferential connectivity to the bursting SUB PC population, this feedback mechanism could act as a precision control mechanism, regulating the amplitude, timing, and duration of excitation to SUB bursting, thereby increasing the accuracy of ripple-driven information transfer from hippocampus to cortex during memory consolidation (Kim and Spruston, 2012; Soltesz and Losonczy, 2018; Cembrowski et al., 2018; Böhm et al., 2015; Nitzan et al., 2020; Mizuseki et al., 2011; Valero and de la Prida, 2018; Johantges et al., 2025; Hanson et al., 2025)

### CALB1+ CA1 PCs support adaptation to a changing environment

Chemogenetic inhibition using CNO during a Barnes maze spatial memory task in a memory consolidation and recall paradigm revealed a role for CALB1+ CA1 PCs in supporting an animal’s adaptive response to a changing environment. In the memory consolidation paradigm (CNO given during training phase), both experimental and control animals showed equivalent learning curves, memory performance, and reductions in time spent in the correct quadrant (44.1% for both groups from STM to LTM). This demonstrates the establishment of a spatial map sufficient to consolidate the location of a goal during a spatial memory task is not dependent on the CALB1+ CA1 PCs alone, and that inhibition of these cells does not impact locomotor abilities (Molinari et al., 1996; Airaksinen et al., 1997; Farré-Castany et al., 2007). Similarly, during the memory recall paradigm (CNO given before testing), both groups displayed equal learning abilities during training. However, once the escape box was removed during the testing phase in the STM and LTM test, control animals could be seen to rapidly reduce time in the correct quadrant (56.9% reduction), while CNO-injected animals only displayed a minimal change in behavior (12% reduction). They remained close to the previous goal location for longer, while control animals pivoted back to renewed exploration. This behavior was observed specifically after the first correct nose poke (Fig: 3B, C, D). The analysis of their cognitive performance suggests that experimental, as well as control group were able to sufficiently develop and improve search strategies required to find the correct target location in the same manner (Fig: 4B1, B2). Overall, our findings extend previous work showing that general dorsal CA1 PC are required for reward encoding (Jarzebowski et al., 2022), and inhibition impairs adaptation to changing reward conditions (Jeong et al., 2018). By specifically targeting the CALB1+ CA1 PCs within the dorsal CA1, we demonstrate that this subpopulation is particularly important for the cognitive flexibility required to update spatial maps, while not being essential for initial learning or memory consolidation.

### Competition of established and novel spatial maps and the role of CALB1+ CA1 PCs in cognitive flexibility

A potential explanation for these findings is the necessary establishment of a new spatial map when reward locations change to capture the updated situation. It then must compete with the previously established spatial map. In our experiments, inhibition of CALB1+ CA1 PCs during the recall phase may have disrupted the formation of a new spatial map representation in which there is no escape box (reward) to be found. While initial map establishment and reward location remained intact, the new representation with fewer active PCs within the dorsal CA1 due to CALB1+ inhibition would be less competitive against the established representation. Additionally, any representation containing intact and firing CALB1+ CA1 PCs might be more effective in the establishment of a spatial map overall, as several studies have found CALB1+ superficial CA1 PCs to support stable place map maintenance and the encoding of global cues (Esparza et al., 2025; Valero et al., 2015; Valero and de la Prida, 2018; Sharif et al., 2020). It must be noted, that this does not fully align with findings by Ding et al. (2022), who observed that CALB1+ PCs in dorsal CA1 showed weaker spatial tuning compared to CALB1-negative deep PCs during juxtacellular recordings combined with optotagging (Ding et al., 2022). This discrepancy could be explained if the CALB1+ CA1 PCs are not encoding a global map, but route vectors towards a goal location (Ormond and O’Keefe, 2022), which would need to be updated upon re-location of the target. Other potentially confounding factors may be that another level of heterogeneity exists within the CALB1+ CA1 PC population of the dorsal CA1. Cavalieri et al. had already noted a third CALB1+ PC population within the deep layer of the CA1 (Cavalieri et al., 2021), and observations of “REM-shifting” CA1 PCs in the deep layer by Mizuseki et al. were later expanded, when a subset of CALB1+ CA1 PCs was found to do the same a few years later (Mizuseki et al., 2011; Gu et al., 2023). It is therefore necessary to remain open to the possibility of further heterogeneity even within this subpopulation of CA1 PCs.

### Significance, implications and future direction

Our findings add further evidence of the functional segregation of the CA1 extends through the hippocampal formation not just along different axes, but also within cellular populations defined by genetic, physiological and developmental properties. It is not just their location within the CA1 that shapes their function and allows the integration and translation of signals as they leave the hippocampal formation to consolidate different types of memories. Cognitive flexibility deficits are hallmarks of aging and neurodegeneration, and progressive CALB1 loss has been observed in Alzheimer’s models (Kook et al., 2014; Palop et al., 2003; Sun et al., 2008). Future studies might study both CA1 sublayers simultaneously, as seen in Esparza et al. (2025) using a Chrna7-Cre mouse line to selectively target deep layer CA1 PCs (Esparza et al., 2025). This will allow us to understand how both sublayers differentially connect to SUB PC populations, and future viral techniques may allow even more selected targeting of cellular subtypes within the dorsal CA1. Furthermore, is it unclear what the underlying mechanisms of spatial learning may be on a structural level at the synapse. Berndt et al. were able to observe an increase in synaptic strength of recently active cells in the superficial layer of the CA1 as an animal traverse a linear track, and additionally saw higher rates of participation and firing of the superficial layer in SWRs (Berndt et al., 2023). We have proposed a role for the CALB1+ CA1 PCs and their distinct inhibitory circuit within the dorsal CA1 to regulate the signal-to-noise ratio as information is processed and transferred to the hippocampal output area. As the bursting SUB PCs are known to propagate SWRs downstream, studying their relationship to LTP by observing changes in presynaptic plasticity, specifically changes in vesicle availability, could be the way towards a deeper understanding of how CALB1+ CA1 PCs gate synaptic strengthening at hippocampal output synapses during spatial map development and updating (Orlando et al., 2021; Nitzan et al., 2020; Böhm et al., 2015; Berndt et al., 2023). By using optogenetic tools to control presynaptic cAMP concentration (Oldani et al., 2020; Wietek et al., 2024) it would be possible to selectively manipulate synaptic strength during a spatial learning tasks, providing a way to test whether presynaptic plasticity at this specific connection gates information transfer during the establishment and updating of a spatial map. By furthering our understanding of specialized cellular subpopulations and how they support the formation and competition of established and novel spatial maps at the cellular substrate level, we may not only explain why it takes days to adapt to our new surroundings after rearranging our office, but also reveal how this adaptive process can fail, and be restored, in neurodegenerative processes.

## ACKNOWLEDGEMENTS

We would like to thank Monika Dopatka, Susanne Rieckmann and Anke Schönherr for their valuable technical and administrative assistance. Funding sources: German Research Foundation: project 327654276 – SFB 1315 (DS), project 184695641 – SFB 958 (DS), project 431572356 (DS), project 415914819 – FOR 3004 (DS); Germany’s Excellence Strategy – Exc-2049-390688087 (NeuroCure to DS and RPS), the European Research Council Horizon 2020 grant 810580 – BrainPlay (DS) and the Federal Ministry of Education and Research project 01GQ1420B – SmartAge (DS).

## Author Contributions

**Conceptualisation:** AV, DS

**Behavioral experimental procedures:** AV, DA

**Rabies tracing:** AV, JT

**Injections:** AV, MD

**In-Vitro recordings:** AV, ASt, ASw, RPS, VM

**Analysis:** AV, DP, DA

**Manuscript presentation:** AV

**Manuscript editing:** AV, DS

**Supervision:** DS

**Funding acquisition:** DS

## Data availability

Datasets available from corresponding author upon request.

## Supplementary Information

**Fig. 6.**
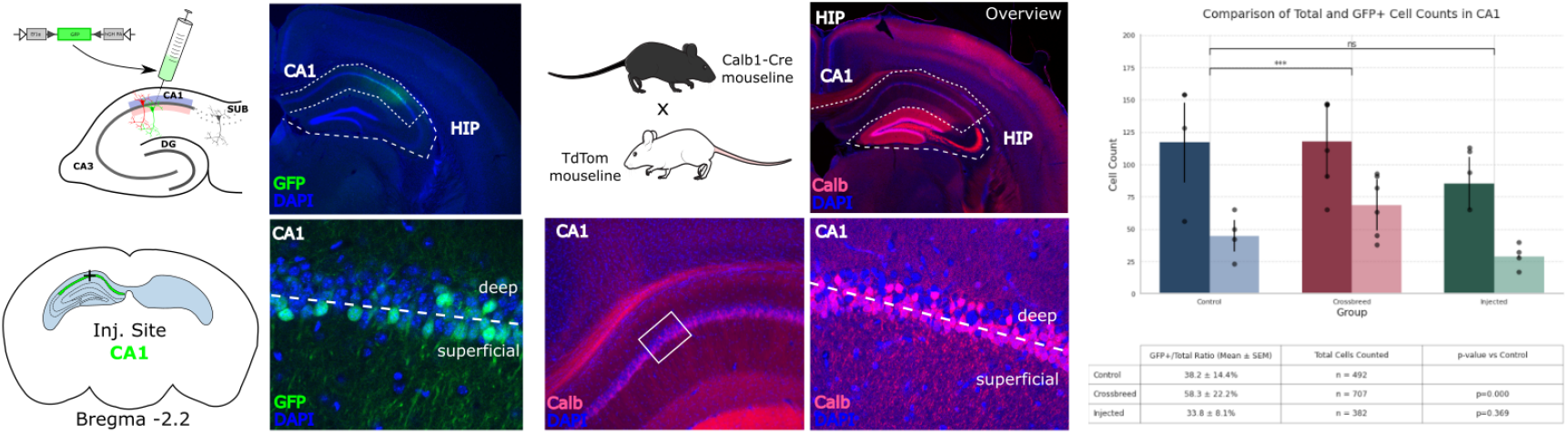
Supplementary Figure: Comparing Cre-dependent injections into CA1 layer of CALB1-Cre animals vs Crossbreeds with reporter line (Ai9) Schematic shows injection location in CALB1-Cre mouseline, overview of the GFP expression in hippocampus (2.5x), zoom in CA1 10x and 20x to see GFP-labeled CA1 PCs in stratum pyramidale. Quantification of labeled cells in superficial CA1 layer for injections vs crossbreeds. GFP-positive cells detected in the deep layer (N(animals) = 2-3 per condition; N(WT-stain) = 492; N(CALB1-CrexTdTom) = 707; N(Cre-dependent injection) = 382).

In these control animals, GFP expression was observed primarily in the superficial layer of CA1, with only a few isolated GFP-positive cells detected in the deep layer (N(animals) = 2-3 per condition; N(WT-stain) = 492; N(CALB1-CrexTdTom) = 707; N(Cre-dependent injection) = 382). These rare deep-layer labeled cells are consistent with reports that a small population of CALB1+ PCs cells reside in the deep pyramidal layer (Cavalieri et al., 2021; Soontornniyomkij et al., 2012), and should therefore be expected to emerge. Overall, viral injections allowed for a far more selective labeling of CA1 sPCs compared to genetic crossbreeding. In CALB1-Cre × Ai9 reporter mice, which constitutively expressed TdTom globally in the brain. These results suggest that while the CALB1-Cre line may include a minority of deep-layer pyramidal cells, injections of Cre-based viral constructs confine the expression of the construct primarily to the superficial sublayer of CA1 more reliably.

Calbindin as a marker of a specific subpopulation of cells along the radial axis becomes even more relevant in the context of a finding by Cavalieri et al. (2021): they found a subpopulation of early born cells in the dPC layer, which also expressed high levels of CALB1, and could be shown to receive less inputs from parvalbumin-positive (PV+) basket cells than expected of cells of the dPC layer (Cavalieri et al., 2021). This finding reveals that it is not soma position alone which defines the CA1 PC subpopulation a cell is defined by – in stark contrast how previous studies have chosen to differentiate between them. This has been observed to happen more often deep layer at the proximal end of the CA1 and in a cross-species studies, a distinct CALB+ band of dPCs was seen in dogs neurogenesis (Slomianka et al., 2011). This differential expression may be driven by transcription factors, active during different time periods of neurogenesis. Overall, these findings suggest that instead of two PC subpopulations along the radial axis, there may be a third outlier group, whose electrophysiological properties, circuit partners and genetic identity may be more defined by birthdate than final soma position. Within this context, targeted genetic tools, such as Cre-dependent viral injections into Cre-mouse lines, become essential for precise work tailored to study a specific PC subpopulation, such as the CALB+ sPCs of the CA1, as cellular identity is defined by more than just its location. However, this approach has limitations: we can only target a limited number of neurons, and our injections were restricted to the dorsal CA1. Additionally, for optogenetic experiments, GFP expression of the viral construct is limited to axonal terminals of ChR2-expressing pyramidal cells, making direct assessment of somatic expression patterns within CA1 challenging. To validate our approach, we conducted control experiments comparing injections to crossbreeds (Supplementary Fig: 6), confirming that injections provide more selective labeling of superficial CA1 PCs.

Results from recordings can be supplemented by histological confirmation of CALB1-content. Because of the densely packed nature of the CA1 pyramidal cell layer, this has proven to be difficult: Valero et al. (2015) attempted to record and stain for CALB1, but based findings on soma location (Valero et al., 2015). Improvements of immunostaining techniques have since also been developed (Merino-Serrais et al., 2020). The relevance of this distinction becomes most apparent in light of the findings by Gu et al. (2023). By selectively optogenetically inhibiting CALB1 sPCs in the CA1, they were able to extend the discovery of “REM-shifting” dPCs (Gu et al., 2023; Mizuseki et al., 2011): Whereas the previous study had found only the dPCs to shift to the theta trough when information is encoded, Gu et al. found subset of CALB1-containing cells doing the same during REM. Additionally, they observed that CALB1-containing sPCs were modulated by ripples during slow-wave sleep, while CALB1-negative dPCs are more active during ripple oscillations. This is the clearest finding that indicates the urgent need for selective study of the CA1 PCs based on genetic markers, and not only on soma location.

**Fig. 7.**
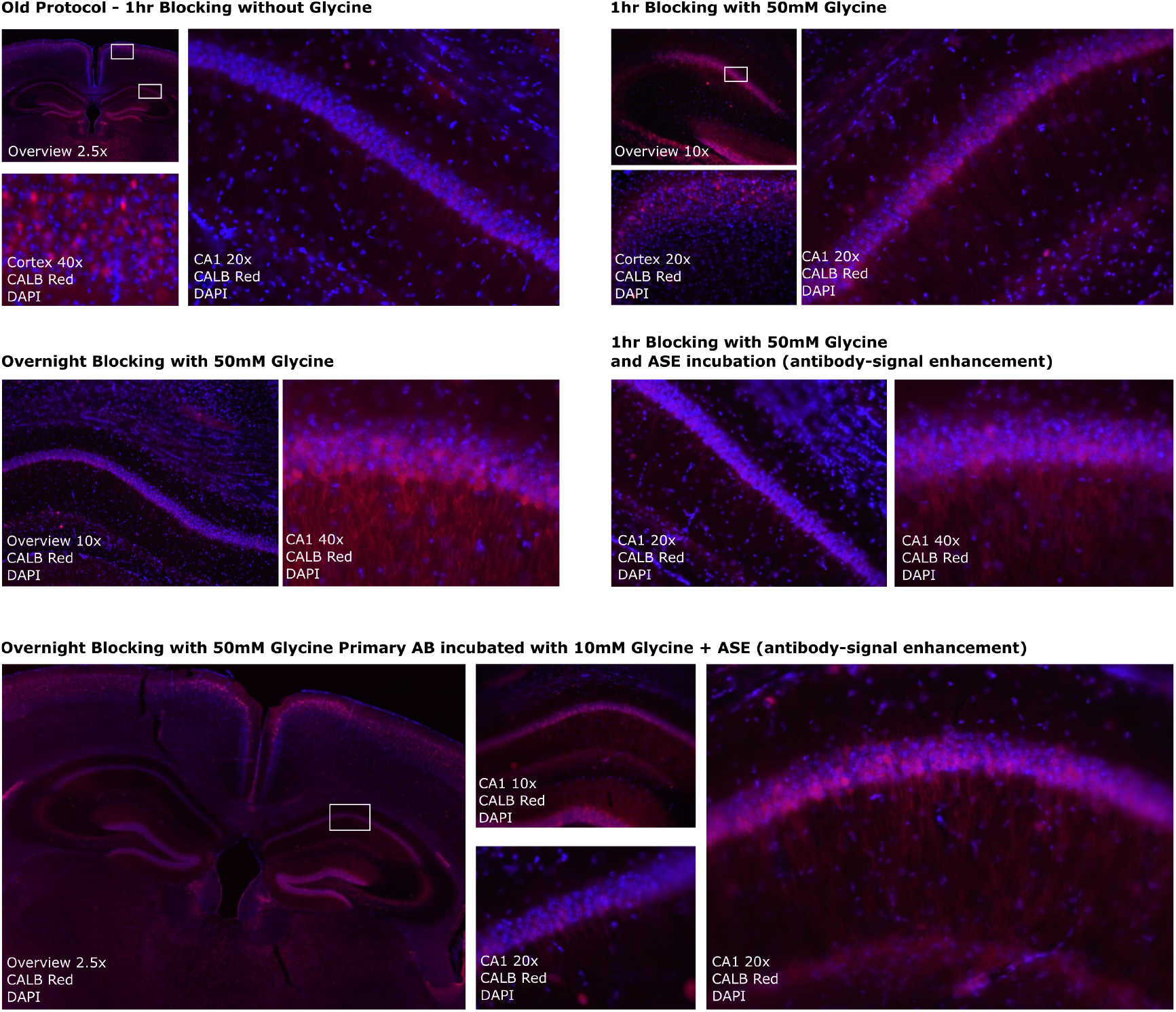
Supplementary Figure: Optimization of Calbindin D28k (CALB1) staining protocol for CA1 PCs. This figure highlights the progressive optimization of CALB1 staining, with the most effective staining observed in panel (C and E). All representative images show stainings of CALB1 in red and DAPI in blue. (A) Original Protocol: Representative images of stainings using the original protocol (Merino-Serrais et al., 2020). Blocking solution without glycine (PBS, 0.1% Triton X-100, 5% NGS). Primary antibody solution contains: PBS, 0.1% Triton X-100, 2.5% NGS, 1:1000 rabbit anti-Calbindin D28k antibody (CB-38a; Swant, Switzerland; 1:2000). Secondary antibody solution contains: PBS, 0.1% 0.1% Triton X-100, 0.5% Tween20, Alexa Fluor anti-rabbit antibody (1:1000). Washing buffer contains: 0.5% Tween20. Shown are an overview image of the mouse brain with focus on the dorsal hippocampus (2.5x), the cortex (40x), and the CA1 pyramidal cell layer (20x). (B) 1hr incubation with 50mM Glycine in blocking buffer: Representative images of stainings using 50mM Glycine in blocking buffer, which is kept on for 1hr. The primary antibody solution also contains 10 mM glycine. Images show overview of dorsal hippocampus (10x), cortex (20x), and CA1 pyramidal cell layer (20x). (C) Overnight incubation with 50mM Glycine in blocking buffer: Blocking performed overnight with 50 mM glycine; the primary antibody solution also contains 10 mM glycine. Shown are an overview of the dorsal hippocampus (10x) and the CA1 pyramidal cell layer (40x). (D) 1hr incubation with 50mM Glycine in blocking buffer and antigen-signal enhancement (ASE) protocol of primary antibody (Gonzalez-Riano et al., 2017). Representative images of stainings using 50 mM glycine in blocking buffer for 1hr; primary antibody solution supplemented with ASE (10 mM glycine, 0.1% H_2_O_2_, 0.05% Tween20, 0.1% Triton X-100). Shown are an overview of the dorsal hippocampus (20x) and the CA1 pyramidal cell layer (40x). (E) Overnight incubation with 50mM Glycine in blocking buffer and ASE protocol: Representative images of stainings using 50 mM glycine in blocking buffer overnight; primary antibody solution according to ASE protocol. Shown are an overview image of the mouse brain with focus on the dorsal hippocampus (2.5x), and the CA1 pyramidal cell layer at 10x and 20x.

We optimized CALB1 immunostaining protocols (Supplementary Fig: 7) to improve visualization and validation of our targeting approach. As described by Merino et al. (2020) and Gonzalez-Riano et al. (2017) immunohistochemical staining procedures can lead not only to variations in staining patterns between different staining conditions, but also inter-experimentally (i.e. differences within the same brain) (Merino-Serrais et al., 2020). To address this, we focused on identifying visible changes in staining intensity and distribution using the antibody rabbit anti-Calbindin D28 K (CB-38a; Swant, Switzerland; 1:2000). Protocol adjustments, including glycine inclusion, extended blocking duration, and antigen signal enhancement (ASE), were evaluated qualitatively rather than quantitatively to determine the conditions that optimize CALB1 labeling in CA1 PCs. Overnight glycine incubation combined with an antigen signal enhancement (ASE) approach used by (Rosas-Arellano et al., 2016) yielded robust, layer-specific labeling of superficial CA1 pyramidal cells. Taken together, these approach supported our validation of the specificity of our optogenetic and anatomical manipulations targeting the CALB1+ CA1 PCs.

**Fig. 8.**
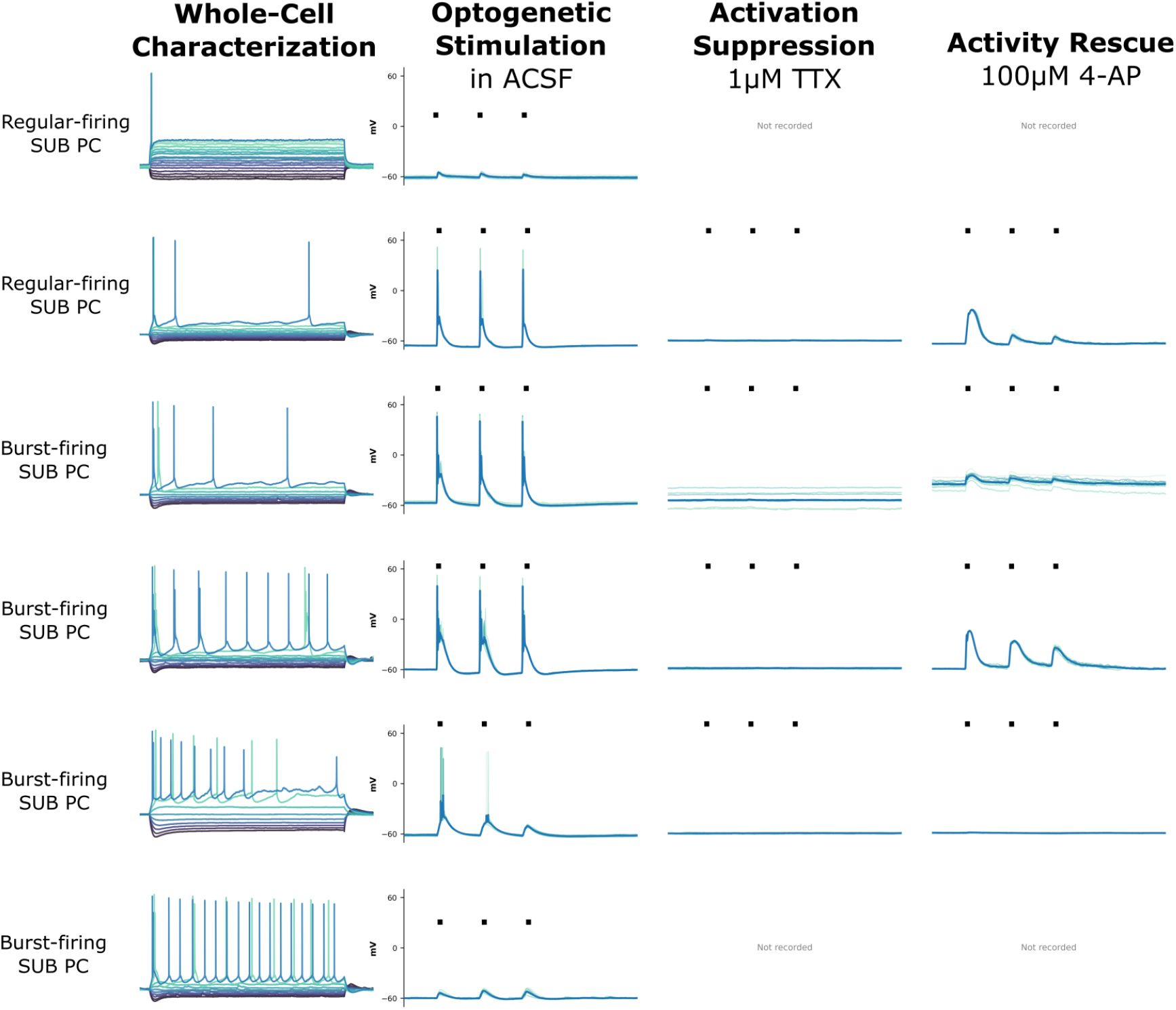
Supplementary Figure: Exemplary Whole-Cell Recording in Multipatch Setting of all 6 Pyramidal Cells. Each row shows SUB PC subtype. Columns from left to right: (1) Cell ID and subtype classification; (2) characterization used for differentiation of cell type (regular-firing, burst-firing SUB PC); (3) light stimulation responses in artificial cerebrospinal fluid (ACSF), these were used to calculate onset delay of responses; (4) activity suppression using 1 *µ*M TTX (traces are absent for cells that did not survive till this experimental condition); and (5) activity rescue after subsequent application of 100 *µ*M 4AP. Recorded fast-spiking interneurons not shown. ■ indicate light activation (10Hz, 10ms).

**Fig. 9.**
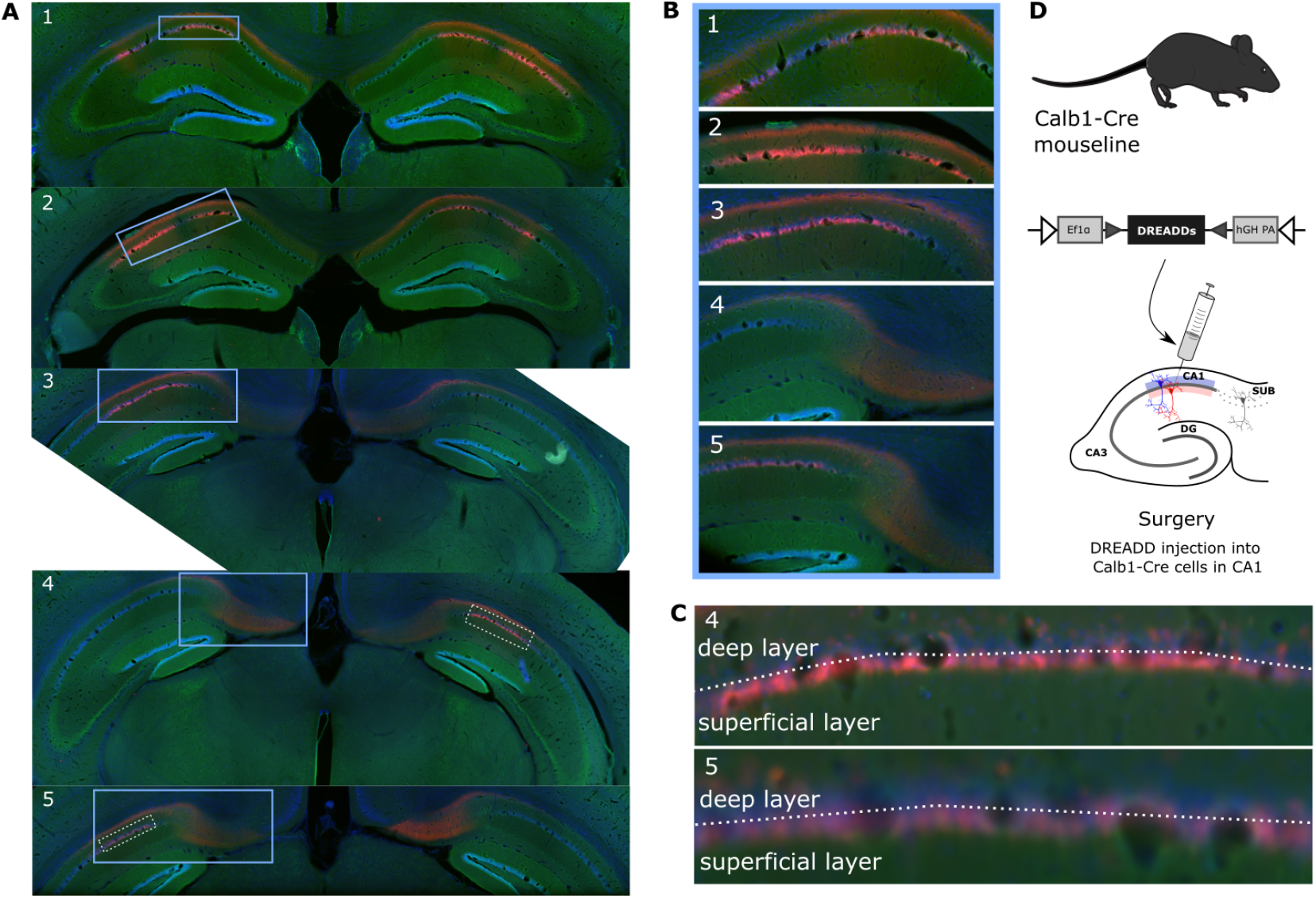
Bilateral DREADD expression in CA1 PCs. Representative histological validation of pAAV-hSyn-DIO-hM4D(Gi)-mCherry injection in CALB1-Cre mouse CA1. (A) Coronal sections showing bilateral mCherry expression (red) in CA1 across the dorsal hippocampus (2.5× magnification, IR-DIC microscopy). (B) Magnification (10×) of selected sections (blue boxes in A) showing detailed bilateral CA1 expression. (C) Magnification (20×) of sections 4 and 5 (dashed boxes in B) revealing mCherry-positive CA1 PCs in superficial sublayer of CA1 PCs. (D) Schematic: Bilateral injection of pAAV-hSyn-DIO-hM4D(Gi)-mCherry into CA1 of CALB1-Cre mice for Cre-dependent DREADD expression in CALB1+ CA1 PCs.

